# Canonical Wnt signaling controls the fate and plasticity of NG2 glia in the healthy and ischemic adult mouse cortex

**DOI:** 10.64898/2025.12.22.696005

**Authors:** Tomas Knotek, Lucie Janeckova, Jan Kriska, Denisa Kirdajova, Alice Foltynova, Jan Kubovciak, Katerina Vecerkova, Makoto Mark Taketo, Michal Kolar, Miroslava Anderova, Vladimir Korinek

## Abstract

NG2 glia, also known as oligodendrocyte precursor cells (OPCs), exhibit unexpected plasticity in the adult brain after injury, yet the molecular cues guiding their fate remain poorly defined. Wnt signaling contributes to tissue responses following ischemic stroke, but its precise role in post-injury glial remodeling is not fully understood. Here, we define how Wnt/β-catenin signaling shapes NG2 glial behavior after focal cerebral ischemia. Using genetic mouse models enabling cell-specific Wnt pathway activation or inhibition, combined with single-cell RNA sequencing, immunohistochemistry, and electrophysiological recording, we disclosed a central role for Wnt signaling in post-injury fate specification.

We identified 12 transcriptionally distinct subpopulations within the oligodendroglial lineage, including a subset with an astrocyte-like transcriptional profile. Wnt signaling strongly influenced the balance between OPC proliferation and differentiation: pathway hyperactivation impaired oligodendrocyte maturation and expanded astrocyte-like NG2-derived cells, likely through concomitant activation of the Notch pathway. Remarkably, Wnt hyperactivation also induced the appearance of cholinergic neuron-like cells derived from NG2-expressing cells exclusively in the somatosensory cortex; these cells generated action potentials and exhibited sodium conductance characteristic of functional neurons.

Together, these findings demonstrate that NG2 glia undergo distinct fate transitions after stroke and that their lineage plasticity is highly sensitive to Wnt pathway dynamics. Targeted fine-tuning of Wnt/β-catenin signaling may enable directed redirection of NG2 glia toward specific reparative outcomes, including neuronal reprogramming, in the injured adult brain.

**TEASER:** Wnt activation promotes neuronal conversion of NG2 glia, while impairing oligodendrocyte maturation.

## INTRODUCTION

Focal cerebral ischemia (FCI), commonly referred to as ischemic stroke, remains one of the leading causes of disability worldwide. Despite extensive research efforts, effective treatment options remain limited. Given the restricted capacity of central nervous system (CNS) neurons to regenerate and repopulate damaged tissue, there is an urgent need to develop clinically relevant therapeutic strategies targeting neuroprotection, immunomodulation, glial scarring, and cellular reprogramming.

Among the glial cells involved in the brain’s response to FCI, cells expressing neuron-glial antigen 2 (NG2), known as NG2 glia, play a particularly important role. They represent the most proliferative cell population outside neurogenic niches in the adult CNS [1] and maintain an even spatial distribution through self-regulatory mechanisms [2]. As oligodendrocyte precursor cells (OPCs), they contribute to learning and memory and are essential for the maintenance of myelin integrity [3]. Beyond the function of OPCs, NG2 glia form functional synapses with neurons and receive both inhibitory and excitatory inputs, which convey the signal to regulate myelination (reviewed in [4]). In addition, NG2 glia are involved in other processes, such as regulation of blood-brain barrier (BBB) thickness and integrity [5, 6], or elimination of thalamocortical presynapses in the visual cortex [7]. Expression of the NG2 protein in the CNS is not restricted to NG2 glia, but also occurs in perivascular cells such as pericytes [8–11]. Importantly, NG2 glia can be reliably distinguished from these perivascular NG2⁺ cells by co-expressing the alpha receptor for platelet-derived growth factor (Pdgfrα), a specific marker of NG2 glia including OPCs (reviewed in [12]).

Following CNS injury, NG2 glia, together with microglia, are among the first cell types to respond to traumatic brain injury. Their response includes proliferation, migration, morphological changes, accumulation near the lesion site, and participation in glial scar formation [1, 13, 14]. A similar role for NG2 glia has also been observed in the context of ischemic stroke [9, 15, 16]. In addition, they probably play a crucial role in ensuring timely wound closure [13]. The differentiation of NG2 glia into oligodendrocytes (OLs) after injury has been demonstrated in several studies [11, 12, 15]. Given the high susceptibility of oligodendrocytes to ischemic damage, NG2 glia are thought to serve as a reserve pool for their replacement (reviewed in [17]).

Interestingly, several studies have shown that the differentiation potential of NG2 glia is not restricted to the OL lineage. *In vitro* experiments have disclosed that these cells can differentiate into astrocytes and in some cases even into neurons. However, *in vivo* lineage tracing studies have yielded conflicting results regarding the neurogenic potential of NG2 glia in the healthy adult brain and following injury (reviewed in [18]). Current evidence suggests that astrogliogenesis from NG2 glia under physiological conditions is largely restricted to prenatal development (reviewed in [12]). Nevertheless, a subset of NG2 glia has been shown to give rise to astrocyte-like cells after injury that express astrocytic markers such as glial fibrillary acidic protein (GFAP) [9–11, 19, 20]. In our recent study, we demonstrated that the gene expression profile of astrocyte-like NG2 glia, analyzed three days after stroke, closely resembles that of neural stem/progenitor cells. Moreover, one month after stroke, we identified NeuN-positive cells derived from NG2-expressing cells in mice [9]. These findings contribute to the ongoing debate regarding the differentiation potential of NG2 glia and their capacity for neurogenesis. We hypothesize that appropriate stimulation could direct NG2 glia toward specific fate outcomes by regulating their proliferation, maintenance, and differentiation, as well as other aspects of their injury response.

One of the key signaling pathways that regulate these processes is the Wnt signaling pathway [21, 22]. Alterations in Wnt signaling have been observed in various pathological conditions, including FCI, and affect multiple brain cell types. Following focal ischemia, Wnt signaling is associated with both neuroprotective and regenerative responses (reviewed in [23, 24]). In one of our previous studies, we reported increased expression of *Wnt7b* and Wnt-responsive genes *Axin2* and *Troy* in NG2 glia in the subventricular zone (SVZ) following FCI [25]. Wnt/β-catenin signaling from NG2 glia has been shown to contribute to BBB integrity and promote angiogenesis after stroke in white matter [26, 27]. Furthermore, Wnt signaling contributes to the regulation of NG2 glia differentiation into mature, myelinating OLs [28].

In this study, we investigated the effects of Wnt signaling on cortical NG2 glia and their progeny following FCI. We employed transgenic mouse models enabling genetic labeling of NG2 glia and targeted modulation of the Wnt/ β-catenin signaling pathway in NG2-expressing cells. Using single-cell RNA sequencing (scRNA-seq), we characterized these cells based on their gene expression profiles. In total, we identified 12 subpopulations within the OL lineage: six subpopulations co-expressing *NG2* and *Pdgfra* genes (*NG2/Pdgfra*⁺ glia), and six corresponding to more differentiated or mature states. Modulation of Wnt signaling markedly shifted the proportions of these subpopulations, and Wnt/β-catenin activation promoted an increased transition of NG2 glia toward an astrocytic fate. This shift may, at least in part, reflect Notch pathway activation triggered by Wnt hyperactivation. Using immunohistochemistry, we identified NeuN-positive cells with neuronal morphology that appeared exclusively in the somatosensory cortex following Wnt hyperactivation and originated from NG2-expressing cells. To assess the functional properties of these cells, we performed patch-clamp electrophysiological recordings, which revealed neuron-like membrane properties and action potential firing, confirming their functional neuronal identity. Finally, immunohistochemical staining of various neuronal markers indicated the cholinergic character of these cells. Our findings underscore the remarkable plasticity of NG2 glia after stroke and highlight canonical Wnt signaling as a key regulator of their lineage fate and regenerative potential.

## RESULTS

### Modulation of the Wnt/β-catenin signaling pathway affects the number and proliferation of NG2-expressing cells and their progeny around the ischemic lesion

To investigate the role of Wnt signaling in the function and differentiation of NG2 glia and their progeny in the healthy and ischemic cerebral cortex of adult mice, we used NG2CreER™ BAC:Rosa26-tdTomato mice that allow inducible, tamoxifen-mediated labeling of NG2-expressing cells. These double transgenic mice harbor the CreER^TM^ cassette, encoding Cre recombinase fused to a modified estrogen receptor (ER) ligand-binding domain that, driven by the *Cspg4* [Cspg4 (chondroitin sulfate proteoglycan 4) is an alternative name for NG2] gene promoter [29]. Thanks to the other genetic modification, tamoxifen administration induces expression of the tandem dimer Tomato (tdTomato) fluorescent protein integrated downstream of the *Rosa26* locus, resulting in red fluorescence in NG2-expressing cells and their progeny; these cells will be hereafter referred to as NG2-tdTom cells. For the purpose of this study, mice from this strain are referred to as controls. To model ischemic stroke, we used middle cerebral artery occlusion (MCAO), in which the MCA is occluded by cauterization. The sham-operated control animals underwent the same surgical procedures, including craniotomy, but without artery occlusion. In this study, the mice that underwent MCAO are referred to as "MCAO mice/animals," while the sham-operated control mice are referred to as "SHAM mice/animals." The time course of the experiment and the methods of analysis are summarized in Figure 1A.

**Figure 1.**
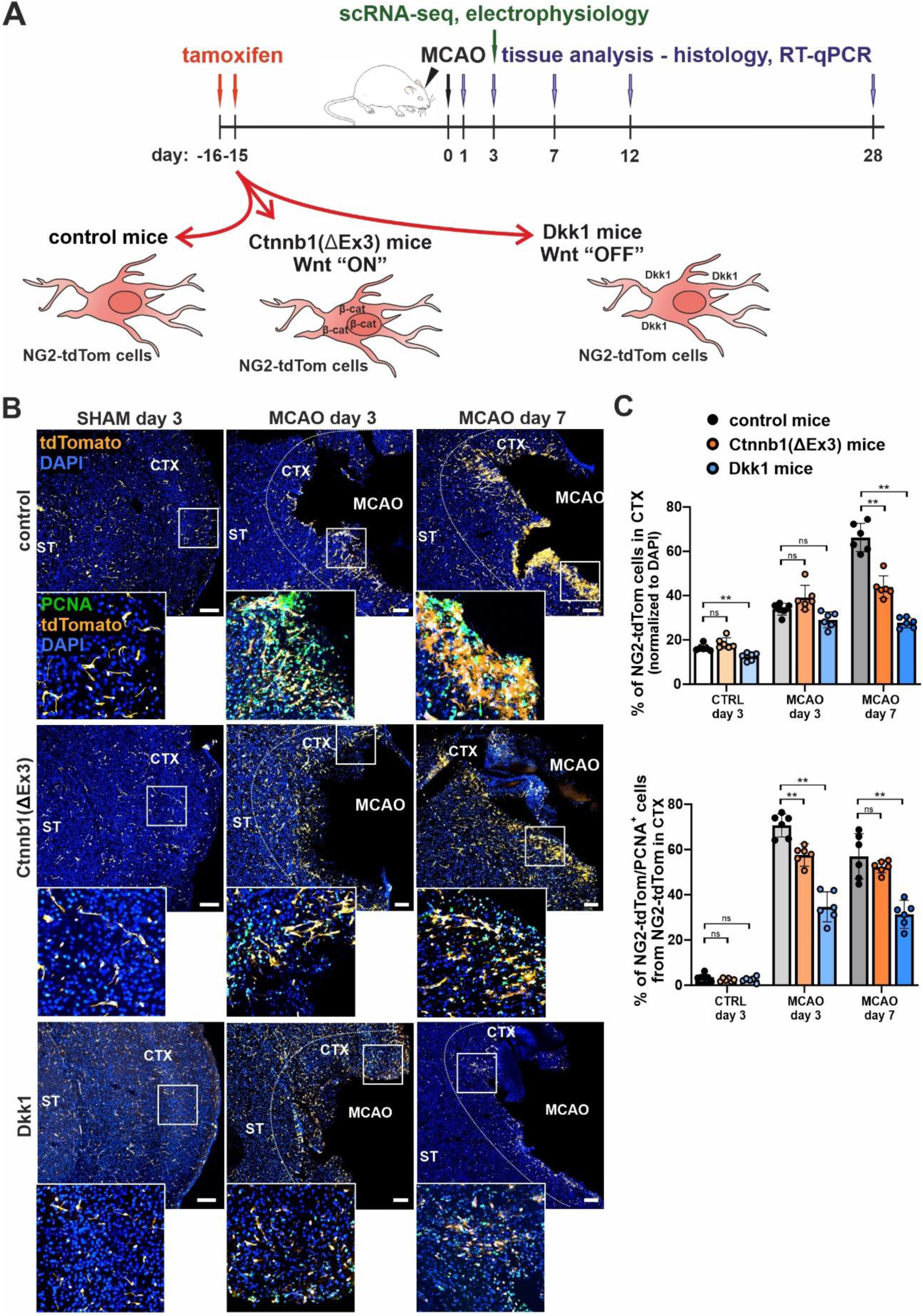
Postischemic hyperactivation or inhibition of Wnt signaling in NG2 glia and their progeny impairs cell proliferation. NG2-expressing cells and their progeny were labeled with tamoxifen-induced tdTomato expression and analyzed in mouse strains with altered Wnt/β-catenin pathway activity. (A) Schematic overview of the experimental setup (top) and the genetic models used to modulate Wnt signaling (bottom). Tamoxifen was administered on two consecutive days (day −16 and −15) to induce tdTomato expression in NG2-expressing cells and their progeny (referred to as NG2-tdTom cells). On day 0, focal cerebral ischemia was induced by permanent middle cerebral artery occlusion (MCAO); sham-operated mice served as controls (SHAM). Brain tissue was harvested 1, 3, 7, 12, and 28 days after surgery and analyzed by single-cell RNA sequencing (scRNA-seq), electrophysiology, histology, and reverse transcription quantitative PCR (RT-qPCR). Mice with unaltered Wnt signaling served as controls and were compared to Ctnnb1(ΔEx3) mice with constitutively active β-catenin (“Wnt ON”) and Dkk1-overexpressing mice with inhibited Wnt signaling (“Wnt OFF”). (B) Representative fluorescence micrographs show NG2-tdTom cells (tdTomato^+^, red signal) in the cerebral cortex (CTX) and striatum (ST) 3 and 7 days after MCAO; the lesion boundary is indicated by a dashed line. The sections were stained with PCNA (green) to visualize proliferating cells and with DAPI (blue) to mark the nuclei. The insets show PCNA^+^ and tdTomato^+^ cells. Scale bar: 150 µm. (C) Quantification of NG2-tdTom cells (top) and NG2-tdTom/PCNA^+^ proliferating cells (bottom) in CTX of SHAM and MCAO animals of the three genotypes. Data are presented as percentage of total DAPI^+^ nuclei (mean ± SEM; n ≥ 6 slices from two mice per group). One-way ANOVA: p < 0.05 (*), p < 0.01 (**), ns: not significant.

Alterations in Wnt signaling are associated with various brain disorders, including Alzheimer’s disease and ischemic stroke (reviewed in [23, 24]). Notably, plasma levels of the Wnt inhibitor Dickkopf-1 (DKK1) are elevated in patients following acute ischemic stroke [30]. We investigated how ischemic stroke affects the mRNA levels of several Wnt/β-catenin–responsive genes that are actively expressed in brain tissue upon modulation of Wnt signaling, namely *Axin2*, naked cuticle 1 (*Nkd1*), specificity protein 5 (*Sp5*), tumor necrosis factor receptor superfamily member 19 (*Tnfrsf19*), and cyclin D1 (*Ccnd1*) [25]. The expression of these genes in NG2-tdTom cells isolated from SHAM mice served as a control. Wnt/β-catenin–responsive genes, except for *Tnfrsf19*, were significantly upregulated three days after MCAO-induced ischemia, as shown in Supplementary Figure S1A and previously reported [25].

Based on our findings that FCI affects Wnt signaling in NG2 glia, we used two transgenic mouse strains with tamoxifen-inducible modulation of the Wnt signaling pathway. To investigate Wnt inhibition, NG2CreER™ BAC:Rosa26-tdTomato mice were crossed with Rosa26-Dkk1 mice [31] to allow expression of Dkk1 (an extracellular Wnt inhibitor) upon activation by tamoxifen. In the resulting progeny (hereafter referred to as the “Dkk1 mice”), Dkk1 can modulate Wnt signaling through autocrine effects within NG2-tdTom cells and through paracrine or juxtacrine effects on neighboring cells. (Hyper)activation of Wnt signaling in NG2-expressing cells was achieved by crossing NG2CreER™ BAC:Rosa26-tdTomato mice with Catnb^lox(ex3)^ mice carrying a floxed exon 3 in the *Ctnnb1* gene encoding β-catenin [32]. Exon 3 contains phosphorylation sites that are critical for targeting β-catenin to the proteasome, such that its deletion stabilizes β-catenin and promotes its accumulation [33]. In the progeny – hereafter referred to as “Ctnnb1(ΔEx3)” mice - these effects lead to sustained β-catenin accumulation in NG2-tdTom cells. Stabilized β-catenin migrates to the nucleus and regulates the expression of canonical Wnt target genes (reviewed in [23, 34]). To validate our transgenic models, we examined the effects of modulating the Wnt signaling pathway in NG2-tdTom cells by RT-qPCR.

In Ctnnb1(ΔEx3) mice, relative mRNA levels of all tested Wnt target genes were elevated compared to controls, reaching stronger statistical significance in MCAO animals. In contrast, Dkk1 mice exhibited reduced expression of these genes; most notably, *Axin2* and *Sp5* mRNA levels were significantly lower in both SHAM and MCAO mice (Supplementary Figure S1B). Additionally, NG2-tdTom cells isolated from Dkk1 mice displayed robust expression of the *Dkk1* transgene, confirming successful induction (Supplementary Figure S1C).

NG2-tdTom cells accumulate around the ischemic lesion both 3 and 7 days after MCAO (Figure 1B), which is consistent with our previous findings [9]. Accordingly, we observed a peak of NG2-tdTom cell proliferation at day 3, which decreased again at day 7 (Figure 1C, top panel). Suppression of Wnt signaling in Dkk1 mice resulted in a marked reduction in NG2-tdTom cell number at day 7 and significantly lower proliferation at both time points. Interestingly, hyperactivation of the Wnt signaling pathway in Ctnnb1(ΔEx3) mice resulted in a similar, albeit smaller, decrease in the NG2-tdTom cell number and proliferation activity (Figure 1C).

These results demonstrate that the Wnt/β-catenin signaling pathway is active in cortical NG2 glia and is enhanced in response to ischemic injury. Disruption of this signaling pathway, whether by inhibition or excessive activation, impairs NG2 glia proliferation. This suggests that precise Wnt regulation is crucial for their proliferation and possibly also for their differentiation and survival after stroke.

### The Wnt/β-catenin pathway modulates the OPC fate by promoting differentiation and impairing terminal myelination

Building on our previous scRNA-seq profiling of NG2-tdTom cells isolated from ischemic or sham-operated cortex [9], we applied the same approach to analyze NG2-tdTom cells from Wnt modulating mouse strains after SHAM or MCAO surgery. Sequencing data were processed using the Seurat package [35]. A pooled dataset of NG2-tdTom cells from all three genotypes [control, Dkk1 and Ctnnb1(ΔEx3)] and both experimental conditions (SHAM and MCAO) was used to identify transcriptionally distinct cell clusters. These clusters were mapped to the major (expected) cell populations, including *Pdgfrb-*expressing perivascular cells, oligodendrocyte transcription factor 1 (*Olig1*)-positive cells of the oligodendroglial lineage (i.e., OPCs and their differentiated progeny), and a smaller population of cells expressing allograft inflammatory factor 1 (*Aif1*) (Figure 2A). Gene Set Enrichment Analysis (GSEA) via the Enrichr platform [36, 37] initially classified these cells as microglia, based on the expression of various colony-stimulating factor receptors (*Csfr*; Supplementary Table S1). Immunofluorescence showed minimal co-localization of tdTomato and Aif1 in both the SHAM and MCAO samples (Supplementary Figure S2A), suggesting that the microglia-like cells in our scRNA-seq dataset were likely to be contamination during fluorescence-activated cell sorting (FACS) isolation rather than a true lineage identity. These cells could alternatively represent CNS-infiltrating macrophages, which have been reported to express NG2 in previous studies [38, 39]. The relative abundance of perivascular cells and microglia/macrophages was the same in SHAM animals in all three genotypes. In contrast, the proportion of OPCs (recognized by their expression of the *Pdgfra* gene) varied depending on the genetic modulation of Wnt signaling (Supplementary Figure S2B).

**Figure 2.**
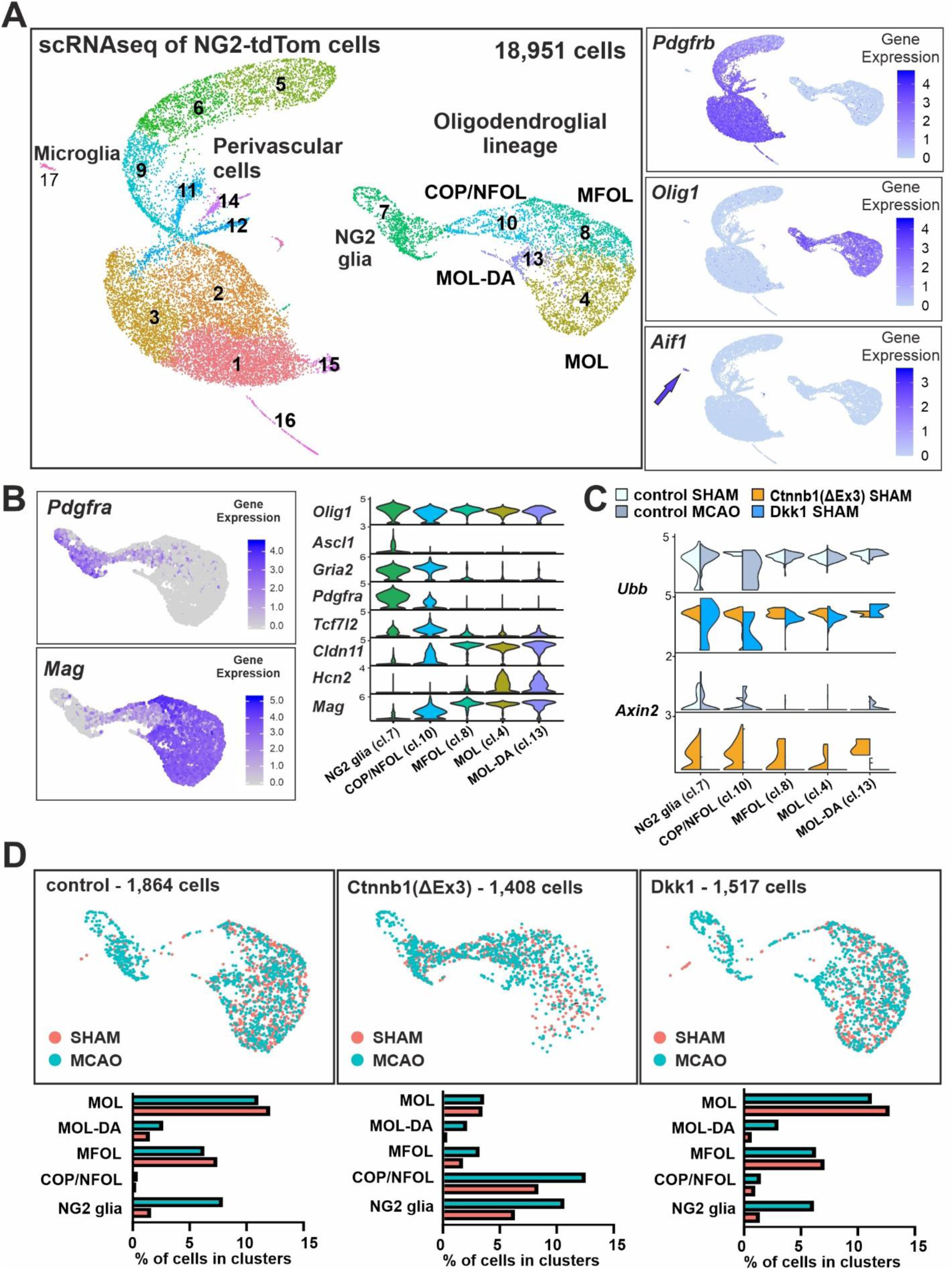
Increased Wnt/β-catenin signaling impairs oligodendrocyte (OL) maturation. Single-cell RNA sequencing (scRNA-seq) was performed on NG2-tdTomato–labeled cells isolated by fluorescence-activated cell sorting (FACS) from the cortex of sham-operated (SHAM) and ischemic (MCAO) mice representing three genotypes: control, Ctnnb1(ΔEx3), and Dkk1. The final integrated dataset comprised 18,951 cells, with an average of 2,639 transcripts per cell. (A) Left, UMAP plot illustrating 17 transcriptionally distinct clusters of NG2-tdTom cells. Based on principal component analysis (PCA) of the integrated dataset, three major lineages were identified: microglia (cluster 17), perivascular cells (clusters 1–3, 5, 6, 9, 11, 12, 14–16), and the oligodendroglial lineage (clusters 4, 7, 8, 10, 13). Right, feature plots showing the expression of representative marker genes for each lineage. The arrow in the bottom plot indicates microglia. (B) Left, feature plots depicting the expression of *Pdgfra* (a marker of early OL lineage cells) and *Mag* (a marker of mature OLs). Right, violin plots showing the distribution of selected stage-specific marker genes across the defined OL lineage clusters: NG2 glia (cluster 7), committed oligodendrocyte precursors/newly formed OLs (COP/NFOL; cluster 10), myelin-forming OLs (MFOL; cluster 8), disease-associated mature OLs (MOL-DA; cluster 13), and fully mature OLs (MOL; cluster 4). (C) Violin plots showing expression of the canonical Wnt pathway target gene *Axin2* across the five OL lineage clusters. The y-axis represents normalized gene expression levels. Ubiquitin B (*Ubb*) expression is shown as a control to illustrate even cell representation across all clusters, as *Ubb* is expected to be ubiquitously expressed. (D) UMAP plots showing OL lineage distribution under SHAM (light red) and MCAO (turquoise) conditions across genotypes. Bar charts beneath each UMAP summarize the relative proportions of cells within each OL lineage cluster, shown as a percentage of the total NG2-tdTom cells analyzed. Abbreviations: *Aif1*, allograft inflammatory factor 1; *Ascl1*, achaete-scute family bHLH transcription factor 1; *Cldn11*, claudin 11; *Gria2*, glutamate ionotropic receptor AMPA type subunit 2; *Hcn2*, hyperpolarization-activated cyclic nucleotide-gated channel 2; *Mag*, myelin-associated glycoprotein; *Olig1*, oligodendrocyte transcription factors 1; *Pdgfra*, platelet-derived growth factor receptor alpha; *Tcf7l2*, transcription factor 7-like 2.

Next, we focused on OL lineage cells because they respond both to modulation of Wnt/β-catenin signaling and ischemic injury [26]. OL lineage cells formed five transcriptionally distinct clusters corresponding to different stages of differentiation (Figure 2A). These clusters were identified based on the expression of stage-specific marker genes associated with OL development and myelination (summarized in [40]): NG2 glia (undifferentiated OPCs), committed OPCs and newly formed OLs (COP/NFOL), myelin-forming OLs (MFOL), mature OLs (MOL), and disease-associated mature OLs (MOL-DA). A complete list of subtype-specific marker genes can be found in Supplementary Table S1.

As in our previous analyses, we found that the expression of glutamate ionotropic receptor AMPA-type subunit 2 (*Gria2*) decreased with OL maturation, whereas the expression of hyperpolarization-activated cyclic nucleotide-gated ion channel 2 (*Hcn2*) increased. The expression of *Pdgfra* was restricted to the early stages of OL differentiation, while expression of the achaete-scute family bHLH transcription factor 1 *(Ascl1*) was specific only to NG2 glia. In contrast, claudin 11 (*Cldn11*) and myelin-associated glycoprotein (*Mag*) were most highly expressed in mature OLs. Of note, expression of transcription factor 7 like 2 (*Tcf7l2*), which encodes T cell transcription factor 4 (Tcf4), the nuclear Wnt signaling mediator, was enriched in cells differentiating along the OL lineage, i.e., in the COP/NFOL cluster. Most strikingly, we identified a population of mature OLs (MOL-DA) containing genes such as kallikrein-related peptidase 6 (*Klk6*), annexin A2 (*Anxa2*), plasmalemma vesicle-associated protein (*Plvap)*, epithelial membrane protein 3 (*Emp3*), and serine (or cysteine) peptidase inhibitor, clade A, member 3N (*Serpina3n*) (Figure 2B, Supplementary Table S1), which have already been associated with CNS pathology, including neurodegenerative diseases [41, 42], and more recently also associated with FCI [43].

To assess the activation state of the Wnt signaling pathway, we examined the expression of the Wnt/β-catenin target gene *Axin2* across the OL lineage (Figure 2C). *Axin2* mRNA was most strongly expressed in NG2 glia and COP/NFOL cells, with levels declining during subsequent OL differentiation. After MCAO, we detected a modest upregulation of *Axin2* in the COP/NFOL and MOL-DA clusters. As expected, *Axin2* expression served as a reliable readout of pathway activation: it was markedly increased in Ctnnb1(ΔEx3) mice and reduced following Wnt inhibition in Dkk1 mice.

Next, we examined how OL subtype distributions varied across the mouse genotypes and treatment conditions (Figure 2D). In MCAO samples, the NG2 glia cluster was proportionally expanded, consistent with the increased number of NG2-tdTom cells observed near the ischemic lesion at days 3 and 7 post-injury (Figure 1B, C). While mature OLs showed a modest decline following ischemia, the disease-associated oligodendrocyte (MOL-DA) subtype was consistently enriched in MCAO tissues, though it remained detectable at low levels in SHAM controls. The OL subtype composition was markedly altered in Ctnnb1(ΔEx3) mice (“Wnt ON”), with a significant reduction in the proportion of myelinating and mature OL clusters (clusters 4, 8) and a corresponding increase in immature populations, particularly NG2 glia and COP/NFOL cells (clusters 7 and 1). This altered distribution pattern persisted after ischemic injury, suggesting that sustained activation of Wnt/β-catenin signaling impairs normal OL lineage maturation.

Although the overall distribution of oligodendroglial lineage clusters appeared largely similar between control and Dkk1 (“Wnt OFF”) mice (Figure 2D), histological analysis revealed a marked reduction in Pdgfrα-positive NG2-tdTom cells in the peri-ischemic cortex of Dkk1 mice at both 3 and 7 days post-injury (Figure 3A, B). This decrease is consistent with the reduced proliferative capacity of NG2-tdTom cells observed in Dkk1 animals (Figure 1C). Differential gene expression (DEG) analysis followed by over-representation analysis (ORA) of NG2 glia from ischemic Dkk1 mice revealed upregulation of several genes associated with cell cycle progression and mitotic regulation (Supplementary Table S2). However, Enrichr analysis [36, 37] identified significant enrichment of phenotypic terms related to abnormal mitotic spindle morphology (MGI Mammalian Phenotype 2024, MP:0009760; adjusted p < 0.0001) and abnormal mitosis (MP:0004046; adjusted p < 0.001). These categories included key regulators of chromosome segregation and mitotic checkpoints, as shown in Figure 3C (top panel). Notably, many NG2 glia-specific genes, including *Cspg4*, *Pdgfra*, *Gria2*, and G protein–coupled receptor 17 (*Gpr17*), a critical G protein-coupled receptor involved in OPC-mediated repair [15], were downregulated in Dkk1 mice. Despite this repression of NG2 glia, remyelination appeared unaffected or even slightly enhanced, as indicated by the presence of MAG-positive OLs in the lesion area (Figure 3D).

**Figure 3.**
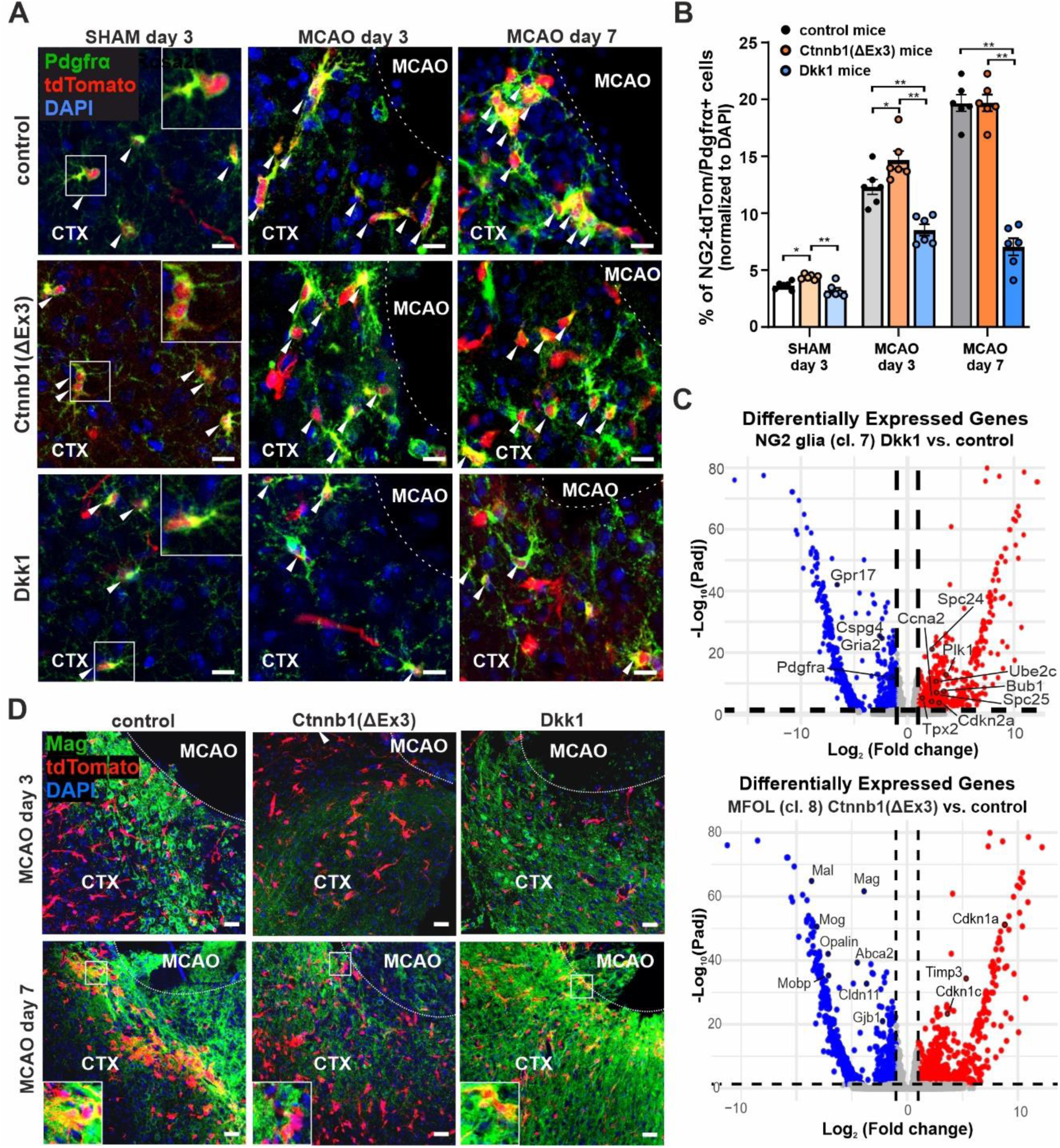
Dysregulated Wnt signaling alters NG2 glia proliferation and myelination after ischemia. (A) Representative fluorescence micrographs of cortical (CTX) sections from SHAM and MCAO mice at 3 and 7 days post-surgery. NG2 glia are identified by co-expression of tdTomato (red) and Pdgfrα (green); nuclei are counterstained with DAPI (blue). Dotted lines indicate the lesion border. Arrowheads highlight tdTomato⁺/Pdgfrα⁺ double-positive cells. Scale bar: 20 μm. (B) Quantification of tdTomato⁺/Pdgfrα⁺ cells in the CTX adjacent to the ischemic lesion at days 3 and 7 post-MCAO across genotypes. Data are expressed as a percentage of total DAPI⁺ nuclei. At least six sections from two animals per group were analyzed (n = 6). Bars represent mean ± SEM. Statistical significance was assessed by repeated-measures ANOVA; *\**p < 0.05; *\*\**p < 0.01. (C) Volcano plots of differentially expressed genes (DEGs) in NG2 glia (cluster 7; top) from Dkk1 mice and MFOLs (cluster 8; bottom) from Ctnnb1(ΔEx3) mice, both compared to controls after MCAO. The x-axis shows log₂ fold change; the y-axis indicates –log₁₀ of the adjusted p-value. Dashed lines denote thresholds for significance (adjusted p < 0.05; |log₂ fold change| ≥ 1). Significantly upregulated (red) and downregulated (blue) genes are highlighted; selected DEGs are labeled. (D) Representative immunofluorescence images of cortical sections at 3 and 7 days post-MCAO. tdTomato (red) marks NG2-lineage cells; Mag (green) labels mature OLs; DAPI (blue) stains nuclei. Scale bar: 50 μm. Abbreviations: *Abca2*, ATP-binding cassette sub-family A member 2; *Bub1*, mitotic checkpoint serine/threonine kinase; *Ccna2*, cyclin A2; *Cdkn1a/c*, cyclin-dependent kinase inhibitor 1A/C; *Cdkn2a*, cyclin-dependent kinase inhibitor 2A; *Gjb1*, gap junction protein, beta 1; *Gpr17*, G protein–coupled receptor 17; *Mal*, myelin and lymphocyte protein; *Mobp*, myelin-associated oligodendrocyte basic protein; *Mog*, myelin oligodendrocyte glycoprotein*; Opalin*, oligodendrocytic myelin paranodal and inner loop protein; *Plk1*, polo-like kinase 1; *Spc24/*25, spindle pole component 24/25; *Timp3*, tissue inhibitor of metalloproteinase 3; *Tpx2*, microtubule nucleation factor; *Ube2c*, ubiquitin conjugating enzyme E2 C.

In contrast, Ctnnb1(ΔEx3) mice (“Wnt-ON”) showed increased numbers of NG2-tdTom–positive cells under both SHAM and MCAO conditions at day 3 (Figure 3A, B). However, MAG immunoreactivity within the ischemic region was markedly reduced at both 3 and 7 days post-injury (Figure 3D), indicating impaired OL maturation and myelin formation. Genes involved in myelination were significantly downregulated in the Ctnnb1(ΔEx3) OL lineage (Supplementary Table S2), particularly in the MFOL subpopulation (Figure 3C, bottom), consistent with the reduced abundance of mature OLs (MOLs) detected in NG2-tdTom cells by scRNA-seq (Figure 2D). Furthermore, the transcriptional profile of Ctnnb1(ΔEx3) samples closely resembled that of Myrf-deficient oligodendrocytes (Enrichr TF Perturbations library: “MYRF KO MOUSE GSE15303 CREEDSID GENE 2306 UP,” adjusted p-value < 0.0001), a gene expression signature associated with failure to initiate myelin gene expression [44]. We also observed upregulation of several genes linked to cell-cycle arrest and apoptosis in the Ctnnb1(ΔEx3) OL lineage, including cyclin-dependent kinase inhibitor 1A (*Cdkn1a*) encoding p21, *Cdkn1c* (p57), and tissue inhibitor of metalloproteinase 3 (*Timp3*; Figure 3C, bottom). These genes are established mediators of OL stress responses and negative regulators of proliferation and differentiation [45–47]. Together, these findings suggest that hyperactivation of canonical Wnt signaling induces a stress-response program that impairs both proliferation and terminal maturation of OLs following ischemic injury.

These data indicate that Wnt/β-catenin signaling has a dual role in the OL lineage. On the one hand, it supports the maintenance and progenitor status of OPCs; on the other, sustained activation impairs terminal maturation and myelination under both physiological and ischemic conditions. Notably, disease-associated mature OLs are present in the ischemic cortex regardless of Wnt status, suggesting that their development is driven more by the ischemic environment than by the Wnt signaling.

### Constitutive Wnt/β-catenin activation drives neuronal-like differentiation of NG2 glia

In our previous work, we identified a subset of NG2-tdTom⁺ cortical cells with the features of astrocytes and neural progenitor characteristics, and we occasionally detected tdTomato⁺ cells expressing the mature neuronal marker NeuN at later stages following FCI, specifically at 28 and 60 days post-injury [9]. Based on this observation, we next investigated whether persistent activation of the Wnt/β-catenin pathway might directly promote neuronal differentiation in this context. This hypothesis was further supported by our earlier findings that Wnt signaling plays a key role in driving neuronal fate from subventricular zone (SVZ)-derived neural stem/progenitor cells [21, 25]. Transcriptomic profiling of NG2 glia – defined here as *NG2⁺Pdgfra⁺* double-positive cells – from Ctnnb1(ΔEx3) mice revealed significant enrichment for gene signatures associated with enhanced neurogenic potential. Notably, DEGs showed strong overlap with transcriptional profiles from forkhead box protein O1 (*Foxo1*) knockout mice (Enrichr, “TF Perturbations Followed by Expression” library; term: FOXO1 KO MOUSE GSE18308 CREEDSID GENE 1168 UP; adjusted p < 0.0001), a model known to exhibit neural stem cell hyperproliferation and brain overgrowth [48]. Additionally, these DEGs were significantly enriched for SRY-box transcription factor 2 (*Sox2*) targets, a transcription factor critical for neurogenesis (Enrichr, “ENCODE and ChEA Consensus TFs from ChIP-X” library; term: Sox2 ChEA; adjusted p < 0.001) [49–51].

To further investigate whether hyperactivation of Wnt signaling promotes neurogenesis from NG2 glia, we examined cortical sections from Ctnnb1(ΔEx3) and control mice three days after SHAM or MCAO surgery –18 days after tamoxifen-induced activation of the Wnt/β-catenin pathway. Notably, we detected NG2-tdTom cells with neuron-like morphology exclusively in Ctnnb1(ΔEx3) brains. These cells were primarily located outside the ischemic lesion, within cortical tissue corresponding anatomically to the somatosensory cortex (based on the mouse brain atlas by [52]). These neuron-like tdTomato-positive cells displayed distinct morphological features, including large somata, a single elongated processes projecting centrifugally toward the cortical surface, and several shorter lateral processes (Figure 4A; Supplementary Figure S3A). Importantly, these cells were observed under both SHAM and MCAO conditions and started appearing 14 days after Wnt pathway activation, suggesting a long-lasting differentiation process (Supplementary Figure S3B).

**Figure 4.**
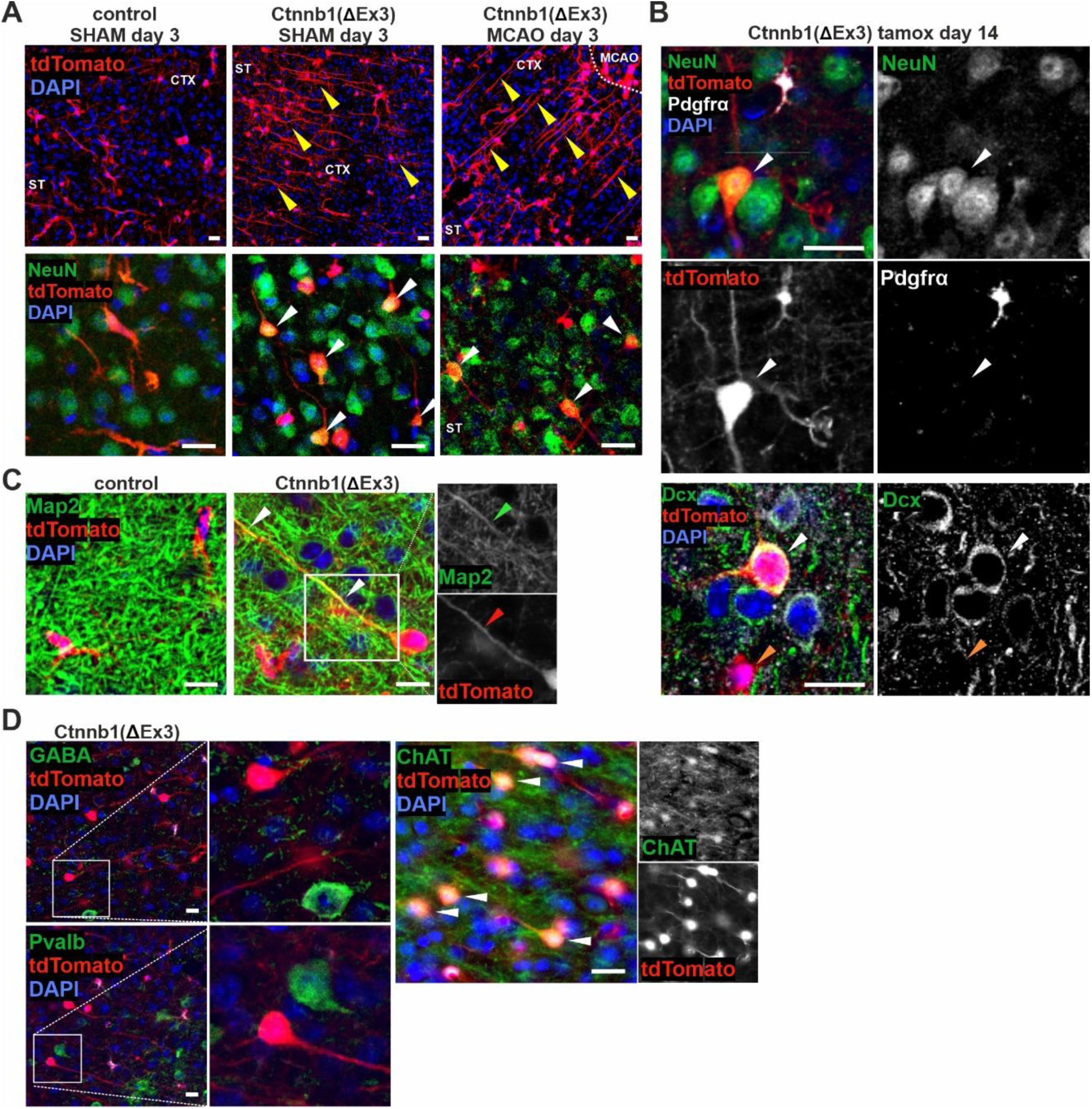
Wnt/β-catenin signaling promotes differentiation of NG2 glia into cholinergic neurons. (A) Representative images of NG2-tdTom cells with long, unbranched protrusions (upper panel, yellow arrowheads) detected in the cortex of both SHAM and MCAO Ctnnb1(ΔEx3) mice, 3 days post-surgery (i.e., 19 days after Wnt/β-catenin pathway activation). A subset of these cells was positive for the neuronal marker NeuN (lower panel, white arrowheads). Scale bar: 30 μm. (B) NeuN-positive NG2-tdTom cells were detectable 14 days after tamoxifen-induced recombination (earlier time points shown in Supplementary Figure S3). These cells exhibited a characteristic long protrusion and lacked Pdgfrα staining (white arrowheads), distinguishing them from smaller, Pdgfrα-positive NG2-tdTom cells with short, branched processes. Some NG2-tdTom cells were positive for doublecortin (Dcx). Scale bar: 30 μm. (C) Long protrusions of NG2-tdTom cells in Ctnnb1(ΔEx3) mice co-localized with microtubule-associated protein 2 (Map2; white arrowheads). In contrast, processes of NG2-tdTom cells in control tissue did not show Map2 staining. Scale bar: 30 μm. (D) Fluorescent images show that NG2-tdTom cells in Ctnnb1(ΔEx3) mice did not express gamma-aminobutyric acid (GABA, upper panel) or parvalbumin (Pvalb, lower panel). In contrast, most neuron-like NG2-tdTom cells were strongly positive for choline acetyltransferase (ChAT), specific for cholinergic neurons. Scale bar: 30 μm.

Immunofluorescence analysis confirmed that these tdTomato⁺ cells expressed NeuN, a marker of postmitotic neurons, but lacked Pdgfrα, indicating a phenotypic transition from glial to neuronal identity. In contrast, canonical OPCs co-expressing tdTomato and Pdgfrα did not stain for NeuN (Figure 4A, B; Supplementary Figure S3B). Additionally, a subset of neuron-like NG2-tdTom cells showed strong immunoreactivity for doublecortin (Dcx), a marker of immature neurons (Figure 4B).

Additional immunostaining confirmed the neuronal identity of these cells. Microtubule-associated protein 2 (Map2), typically localized in neuronal processes [53], was detected in the protrusions of NG2-tdTom neuron-like cells (Figure 4C). To further characterize their neuronal phenotype, we performed immunostaining for the neurotransmitter gamma-aminobutyric acid (GABA) and the calcium-binding protein parvalbumin (Pvalb), two markers of GABAergic neurons commonly found in neurons derived from reprogrammed glia *in vivo* [54]. However, we observed no co-localization of these markers with NG2-tdTom cells. In contrast, choline acetyltransferase (ChAT), a marker of cholinergic neurons, was strongly expressed in a subset of NG2-tdTom neuron-like cells (Figure 4D), suggesting acquisition of cholinergic neuronal identity.

Given the presence of neuron-like NG2-tdTom cells in both SHAM and MCAO Ctnnb1(ΔEx3) mice (as identified histologically), we sought to understand why these cells were not captured in the scRNA-seq dataset, particularly those expressing neuronal genes such as RNA-binding fox-1 homolog 3 (*Rbfox3*; the gene encoding NeuN). Noting the spatial separation of these cells from the ischemic core, we attempted using FACS to isolate Pdgfrα-negative NG2-tdTom cells specifically from this region for quantitative PCR analysis. However, we were unable to detect transcripts for neuronal genes, including calbindin 2 (*Calb2*), *Rbfox3*, or synaptophysin (*Syp*), regardless of Wnt pathway status. Given that the neuron-like cells observed in the tissue clearly possessed NeuN and Map2, we hypothesize that mature neuron-like NG2-derived cells may be especially vulnerable to damage during dissociation and sorting, as we have previously observed with mature neurons isolated from the SVZ and striatum using the same protocol [25].

To address this limitation, we performed functional analysis of NG2-tdTom cells in freshly isolated brain slices. Using whole-cell patch-clamp recordings, we evaluated 53 NG2-tdTom cells from Ctnnb1(ΔEx3) mice. The vast majority (n = 50) showed fast-activating sodium (Na^⁺^) currents in response to depolarization, and 30 cells generated action potential (AP)-like activity in current-clamp recordings (Figure 5A). Both Na^⁺^ currents and AP-like activity were abolished by 1 µM tetrodotoxin, confirming the involvement of voltage-gated Na⁺ channels (Figure 5B). Intracellular labeling with Alexa Fluor 488 contained in the solution confirmed that the recorded cells were neuron-like NG2-tdTom cells with long processes (Figure 5C).

**Figure 5.**
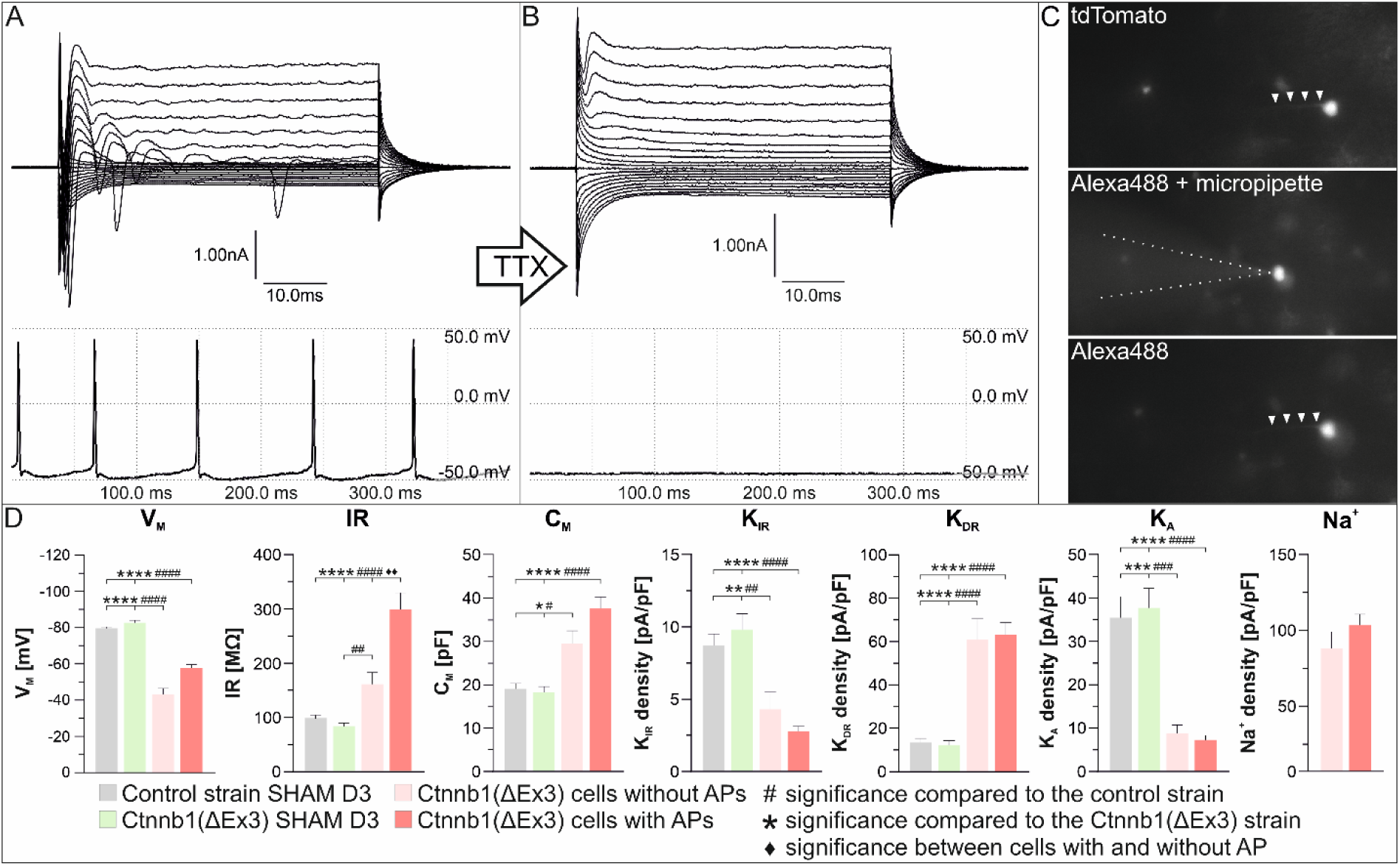
Electrophysiological characterization of NG2-tdTom cells with neuron-like morphology in Ctnnb1(ΔEx3) mice. (A) Representative patch-clamp recordings from an NG2-tdTom-positive cell with a long protrusion. Top: voltage-clamp recording with stepwise depolarization from -160 mV to +40 mV in 10 mV increments, evoking inward Na⁺ currents. Bottom: current-clamp recording showing spiking activity resembling neuronal action potentials. (B) Recordings from the same cell after application of 1 µM tetrodotoxin (TTX), showing complete blockade of voltage-gated Na⁺ currents and action potential-like activity. (C) Fluorescence micrographs of the recorded cell, demonstrating tdTomato expression (red) and Alexa Fluor 488 dye (green) introduced during patch-clamp recording. Arrowheads indicate the elongated protrusion characteristic of neuron-like morphology. (D) Quantitative comparison of electrophysiological parameters between cortical NG2 glia and neuron-like NG2-tdTom cells from Ctnnb1(ΔEx3) mice and control cells. Parameters include resting membrane potential (V_M_), input resistance (IR), membrane capacitance (C_M_), and current densities for Na⁺ currents and K⁺ channel subtypes: inwardly rectifying (K_IR_), delayed rectifier (K_DR_), and A-type (K_A_) K⁺ currents. AP, action potential.

Next, we compared electrophysiological parameters of neuron-like NG2-tdTom cells from non-operated Ctnnb1(ΔEx3) mice, cortical NG2-tdTom glia from SHAM Ctnnb1(ΔEx3) mice, and cortical NG2-tdTom glia from the SHAM control mice, following the analytical framework established in our previous study [9], with the exception of membrane capacitance, which was assessed from area-under-curve of a current elicited by a small hyperpolarizing pulse, as described in the methods section. Parameters included resting membrane potential (V_M_), input resistance (IR), membrane capacitance (C_M_), and densities of potassium (K^+^) currents (K_IR_ for inwardly rectified K^+^ currents, K_DR_ for delayed rectified K^+^ currents, K_A_ for the A-type of K^+^ currents) and Na^+^ currents. Neuron-like NG2-tdTom cells displayed significantly more depolarized V_M_ and elevated IR compared to typical cortical NG2 glia (Figure 5D), suggesting greater excitability [55, 56]. Cells with neuron-like morphology also showed increased C_M_ consistent with larger membrane surface area; this increase was more pronounced in the cells with AP-like activity (Figure 5D, Supplementary Table S3). We observed a reduction in K_IR_ density and an increase in K_DR_ density in neuron-like NG2-tdTom cells, indicating a shift toward a more “outwardly rectified” electrophysiological profile [25]. Additionally, we noticed a significantly reduced density of K_A_ currents. Notably, while none of the NG2 glia recorded in non-ischemic animals (n = 46 for control and n = 37 for Ctnnb1(ΔEx3) NG2-tdTom cells) exhibited Na^+^ currents, nearly all neuron-like cells did. In contrast, Na^+^ currents were sparsely observed in cortical NG2 glia after MCAO (n = 2/40 in the control tissue, n = 0/19 in the Ctnnb1(ΔEx3) tissue, n = 7/29 in the Dkk1 mice). As previously reported [57], IR was also increased after ischemia, but this effect was present only in the control and the Dkk1 strain (Supplementary Table S3).

Together, these results show that activation of the Wnt/β-catenin signaling pathway induces a subset of NG2-tdTom cells (most likely NG2 glia) to acquire a neuron-like phenotype. These cells express neuronal markers and exhibit intrinsic membrane properties, ion channel profiles, and excitability consistent with neurons. Importantly, these neuron-like cells were localized specifically in the somatosensory cortex and observed in both healthy and ischemic conditions in Ctnnb1(ΔEx3) mice.

### Wnt/β-catenin signaling promotes plasticity and preserves progenitor-like features in NG2 glia subpopulations

Next, we sought to better understand the plasticity of NG2 glia following ischemic injury and the extent to which this plasticity is influenced by Wnt/β-catenin signaling. To increase resolution and eliminate confounding signals, we excluded perivascular cells, microglia, and macrophages from the scRNAseq dataset analysis and performed subclustering of cells within the OL lineage. This targeted approach enabled high-resolution characterization of the OL lineage and revealed twelve transcriptionally distinct subpopulations (Figure 6A), refining the original five OL lineage clusters. Cluster identities were defined based on specific gene expression signatures (Supplementary Table S4), supported by GSEA, cell type annotations from the PanglaoDB database [58], and previous single-cell transcriptomic studies of the OL lineage [40].

**Figure 6.**
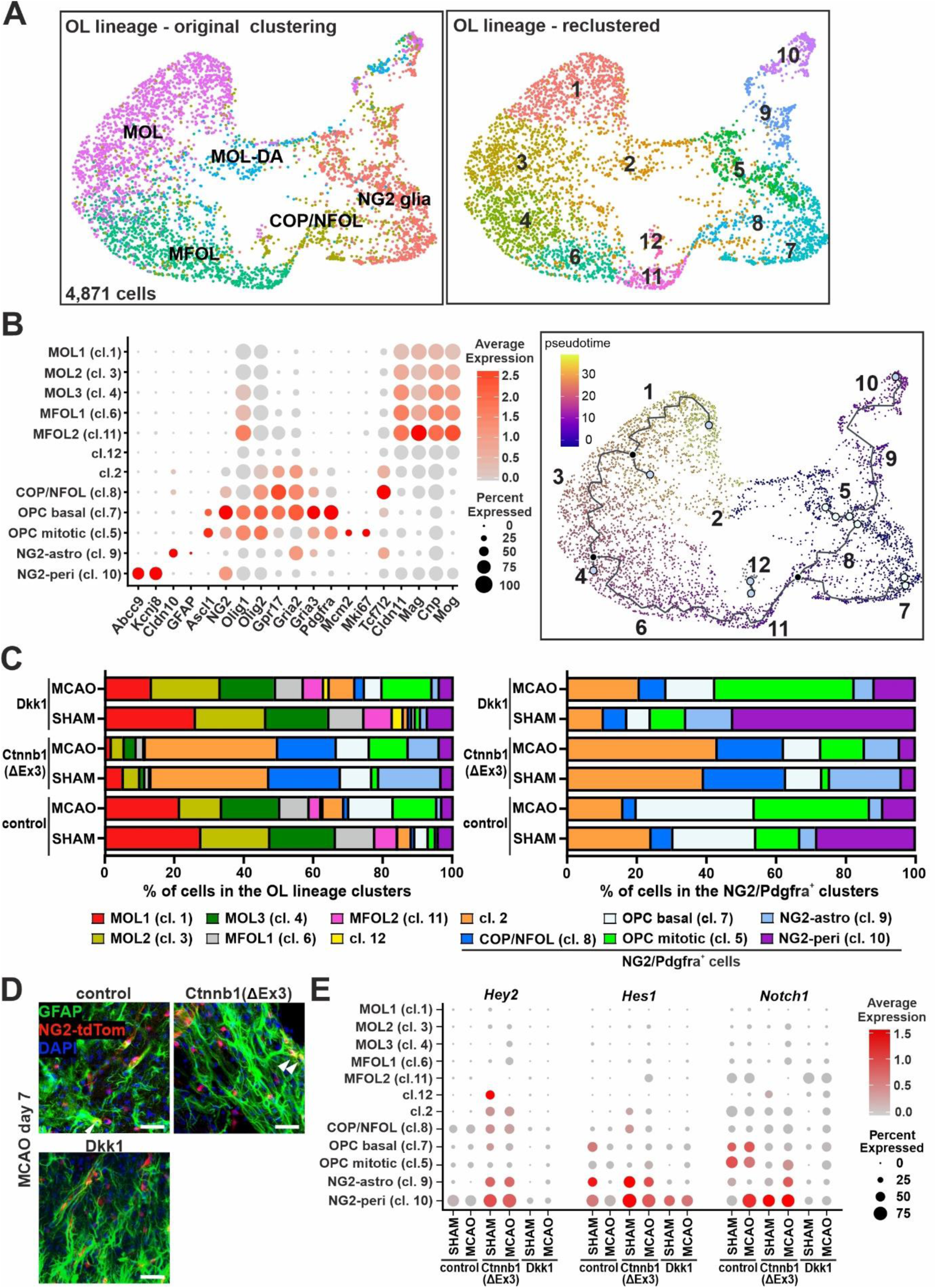
Wnt/β-catenin signaling regulates NG2 glia heterogeneity and cell fate decisions in the OL lineage. Single-cell transcriptomic profiling of NG2-tdTomato⁺ cells from SHAM and MCAO cortices revealed a spectrum of transcriptionally distinct oligodendrocyte lineage (OL) subpopulations, modulated by Wnt signaling activity. (A) UMAP plots show the original clustering into five major OL lineage compartments (left) and the refined re-clustering into twelve transcriptionally distinct subtypes (right), based on exclusion of non-OL cells and higher-resolution analysis. A total of 4,781 OL lineage cells were analyzed. Clusters are labeled by numbers (descending by cell count) and annotated by lineage identity. (B) Left, dot plot shows the expression of key marker genes across the twelve clusters, reflecting lineage identity. Genes mark astrocytes (*Cldn10*, *Gfap*), pericytes (*Abcc9*, *Kcnj8*), OPCs (*Ascl1*, *Olig1/2*, *Gpr17*, *Gria2/3*, *Pdgfra*), proliferating cells (*Mcm*2, *Mki67*), and differentiating or mature OLs (*Tcf7l2*, *Cldn11*, *Mag*, *Cnp*, *Mog*). Dot size represents the percentage of expressing cells within each cluster; color scale indicates mean expression. Several NG2 glial clusters co-express markers of alternative lineages (astrocytic or perivascular), suggesting partial lineage priming or transcriptomic convergence. Right, pseudotime trajectory reconstruction of OL lineage cells using Monocle3 reveals the inferred developmental progression from early precursors (dark purple) to terminally differentiated states (yellow). The black trajectory line represents the learned principal graph; filled circles indicate branching points, and open circles mark trajectory termini. (C) Stacked bar plots show the relative abundance of OL lineage subpopulations under different experimental conditions [SHAM vs. MCAO in control, Dkk1, and Ctnnb1(ΔEx3) mice]. Left, distribution of all twelve OL lineage clusters. Right, focused analysis of NG2 glia (NG2/Pdgfra⁺ cells only), showing Wnt pathway effects on specific subpopulations. (D) Fluorescent micrographs showing GFAP+ astrocytes (green signal) and NG2-tdTom cells (red signal) within the ischemic area 7 days after MCAO. Double-positive cells are marked by white arrowhead. Scale bar: 30 μm. (E) Dot plot showing the expression of Notch signaling-responsive genes (*Hey2*, *Hes1*) and the receptor *Notch1*. Dot size indicates the proportion of cells expressing each gene within a given cluster; color intensity reflects the average expression level. Abbreviations: *Abcc9*, ATP-binding cassette subfamily C member 9; *Cnp*, 2′,3′-cyclic-nucleotide 3′-phosphodiesterase; *Gfap*, glial fibrillary acidic protein; *Hes1*, hairy and enhancer of split 1; *Hey2*, hairy/enhancer of split-related with YRPW motif 2; *Kcnj8*, potassium inwardly rectifying channel subfamily J member 8; *Mcm2*, minichromosome maintenance complex component 2; *Mki67*, proliferation marker Ki-67; *Notch1*, Notch receptor 1.

Cell type-specific gene expression and differentiation pseudotime trajectory is shown in Figure 6B. Clusters 2, 5, and 7–10 expressed canonical OPC markers such as *NG2* (*Cspg4*), *Pdgfra*, and *Gria3*, consistent with their classification as NG2 glia [10]. Within these clusters, however, we identified a transcriptionally distinct subpopulation expressing pericyte-specific genes, including ATP-binding cassette sub-family C member 9 (*Abcc9*) and potassium inwardly rectifying channel subfamily J member 8 (*Kcnj8*), as previously reported [59]. We designated this cluster “NG2-peri” (cluster 10), suggesting a potential perivascular identity within the broader NG2 glial pool.

Another unique subpopulation, designated as “NG2-astro” (cluster 9), was characterized by elevated expression of astrocytic markers, including *Cldn10* and *Gfap* [60], as detailed in Supplementary Table S4. Notably, we previously described a subset of NG2 glia that exhibited gene expression patterns resembling those of astrocytes and neural progenitor cells (NPCs) [9]. These astro/NPC-like cells co-expressed progenitor markers such as *Ascl1* [61], *Gpr17*, and cell cycle-associated genes including minichromosome maintenance complex component 2 (*Mcm2*) and marker of proliferation Ki-67 (*Mki67*). However, the current NG2-astro cluster lacked expression of these proliferative and progenitor markers.

In contrast, these genes were enriched in cluster 5, which also expressed core OPC markers such as *Pdgfra*, *Olig1/2*, and *Gria3* (Figure 6B, left). This cluster, termed “OPC-mitotic,” represents a highly proliferative OPC subset, as further confirmed by cell cycle phase analysis (Supplementary Figure S4).

Importantly, this population formed the starting point of the pseudotime trajectory, a computational inference of the dynamic progression of OL lineage cells through differentiation (Figure 6B, right).

One branch of the pseudotime trajectory [62] extended toward the NG2-astro cluster and then into the slightly more differentiated NG2-peri population, suggesting a potential lineage relationship between these two transcriptionally related subtypes. The second trajectory branch progressed toward a population of committed OPCs and newly formed OLs, designated as COP/NFOL (cluster 8). This cluster was characterized by high expression of *Tcf7l2*, a key transcription factor that drives progression along the OL lineage [10, 11, 63].

In addition to the OPC-mitotic cluster (cluster 5), which showed strong expression of cell cycle–associated genes and marked the proliferative origin of this trajectory, a second origin was identified in cluster 7 ("OPC-basal"). These cells expressed core OPC markers (*Pdgfra*, *Olig1/2*, *Gria3*) along with *Ascl1* and *Gpr17*, but lacked transcripts associated with active proliferation, indicating a quiescent or primed state (Figure 6B, right). Together, clusters 5 and 7 likely represent the same OPC subtype in different functional states, corresponding to actively cycling and non-cycling phases of the cell cycle.

Cluster 2 displayed a “hybrid” transcriptional profile, combining characteristics of both OPCs and more differentiated OLs. It partially overlapped with the MOL-DA population identified in the initial clustering (Figure 6A), but also included cells previously classified as COP/NFOL. Notably, cluster 2 did not align with either major branch of the pseudotime trajectory (Figure 6B, right), suggesting it may represent a non-canonical or arrested differentiation state. This cluster was particularly enriched in Ctnnb1(ΔEx3) mice, coinciding with the expansion of the COP population (Figure 6C). Together, these findings indicate that cluster 2 consists of a heterogeneous population of ischemia-induced MOL-DA cells and OLs whose maturation is stalled by sustained Wnt/β-catenin signaling.

In addition to the *Pdgfra⁺*OPC subtypes, five clusters represented progression along the OL maturation axis (Figure 6B, right). This trajectory extended from committed OPCs and newly formed OLs (COP/NFOL; cluster 8) through two intermediate stages of myelin-forming OLs (MFOL1 and MFOL2; clusters 6 and 11), culminating in three distinct subtypes of mature OLs (MOL1–3; clusters 1, 3, and 4). Expression of key myelin-related genes, such as *Mag*, 2′,3′-cyclic nucleotide 3′-phosphodiesterase (*Cnp*), and myelin oligodendrocyte glycoprotein (*Mog*), peaked during the pre-myelinating MFOL2 stage and gradually declined with terminal differentiation. At the same time, expression of canonical OL lineage genes (e.g., *Olig1/2*, *Gria2*) progressively decreased across MFOL1 and MOL clusters (Figure 6B, left).

Cluster 12, a low-abundance population enriched mainly in Dkk1 mice, lacked expression of typical OPC or OL markers and did not integrate into the reconstructed pseudotime trajectories (Figure 6B). These cells likely represent dysfunctional or aberrant OLs resulting from disrupted Wnt signaling.

Among cells isolated from the cortex of Ctnnb1(ΔEx3) mice, the vast majority (>80%) of OL lineage cells retained expression of *NG2* and *Pdgfra*, indicating a block in lineage progression. Notably, the NG2-astro cluster was markedly enriched in these mice – even under SHAM conditions – suggesting that Wnt/β-catenin pathway activation promotes an alternative astrocyte-like fate within the NG2 glial population (Figure 6C). In addition, enhanced GFAP staining was observed in the ischemic area of Ctnnb1(ΔEx3) mice, including the presence of tdTomato/GFAP double-positive cells (Figure 6D).

In contrast, SHAM Dkk1 mice showed an increased representation of NG2-peri cells and a reduction in mitotic OPCs, indicating that Wnt inhibition limits OPC proliferation and shifts the fate toward a pericyte-like phenotype (Figure 6C, right). Interestingly, ischemic injury partially reversed this effect in Dkk1 mice, restoring the abundance of proliferating OPCs (Figure 6C). However, this shift appeared exaggerated in the scRNA-seq data, as the injury-induced reduction in other cell types led to an apparent increase in the relative proportion of proliferating OPCs, even though the number of dividing NG2-tdTom cells remained lower than in controls (Figure 1B, C). Finally, both Wnt pathway activation and inhibition resulted in a marked reduction of basal/quiescent OPCs across both SHAM and MCAO conditions (Figure 6C, left), suggesting that proper Wnt signaling levels are essential for maintaining this progenitor pool.

Because neuronal cells were not captured in the NG2-tdTom scRNA-seq dataset, we instead aimed to identify NG2 glial subpopulations with potential neurogenic capacity. A closer analysis of the cellular clusters – specifically, assessing the effect of individual genotypes on differences between cells assigned to each cluster – showed the most pronounced expansion in Ctnnb1 (ΔEx3) mice in three clusters: cluster 2 (MOL-DA cells), cluster 8 (COP/NFOL cells), and cluster 9 (NG2-astro). This expansion occurred at the expense of clusters containing more differentiated cells (clusters 1, 3, 4, 6, and 11; Figure 6C).

Analysis of gene expression in cluster 2 revealed features of a heterogeneous cell population. Enrichr ORA of DEGs upregulated in cluster 2 of Ctnnb1(ΔEx3) mice following MCAO, compared with controls (Supplementary Table S5), showed enrichment for genes associated with neural crest differentiation (WikiPathways 2024 Mouse; WP2074; adjusted p < 0.05), including genes responsive to the Wnt and Notch pathways. Moreover, DEGs upregulated in cluster 8 (COP/NFOL) cells from Ctnnb1(ΔEx3) mice exhibited significant overlap with markers of quiescent neural stem cells (NSCs), OPCs, and embryonic or adult astrocytes in both mouse and human datasets (CellMarker 2024; all adjusted p < 0.0001). Conversely, genes downregulated in Ctnnb1(ΔEx3) COP/NFOL cells were enriched for OL lineage signatures in both species (CellMarker 2024; adjusted p < 0.0001), representing broad OL-associated transcriptional programs rather than markers of terminally differentiated, myelinating OLs. These expression patterns were observed in both SHAM and MCAO conditions, with ischemia-specific results shown in Supplementary Table S5.

Cluster 9 (NG2-astro), another population expanded in Ctnnb1(ΔEx3) mice, showed a marked increase predominantly in SHAM animals. DEGs upregulated in this cluster in Ctnnb1(ΔEx3) mice were strongly enriched for embryonic astrocyte and OPC markers across mouse and human datasets (CellMarker 2024; adjusted p < 0.00001). In contrast, genes upregulated in the same cluster in control mice were enriched for markers of mature astrocytes from multiple regions, including mouse striatum, spinal cord, and cerebrospinal fluid (CellMarker 2024; adjusted p < 0.01–0.0001).

The Notch signaling pathway plays a central role in cell fate determination in the CNS and regulates several critical processes during glial development, including maintaining NG2 cells in an undifferentiated state (reviewed in [64]). Wnt signaling has been shown to activate the Notch pathway or key components of its signaling machinery in the CNS, either through direct transcriptional regulation or indirectly [65, 66]. Consistent with this, we observed increased expression of the Notch-responsive genes hairy and enhancer of split 1 (*Hes1*) and hairy/enhancer of split-related with YRPW motif 2 (*Hey2)* in Ctnnb1(Ex3) mice, predominantly in clusters 9 (NG2-astro) and 10 (NG2-peri) in both SHAM and MCAO samples (Figure 6E). Additionally, *Notch1* mRNA was primarily detected in NG2-peri cells in Ctnnb1(ΔEx3) mice under both SHAM and MCAO conditions, as well as in control mice after ischemia.

Collectively, our findings highlight the remarkable plasticity and heterogeneity of NG2 glia. Although their primary fate is differentiation into OLs, we identified NG2-tdTom cells with transcriptional profiles resembling astrocytes and perivascular cells; however, transcriptomic similarity alone does not constitute definitive evidence of lineage conversion. Activation of the Wnt/β-catenin pathway markedly increased the abundance of the NG2 astrocyte-like subpopulation and disrupted normal OL lineage progression. Sustained Wnt signaling interfered with terminal OL maturation, maintained NG2 glia in a plastic, progenitor-like state, and counteracted astrocytic maturation in NG2-astro cells, resulting in lineage ambiguity and delayed fate commitment. These findings underscore the importance of precise temporal control of Wnt signaling, as transient activity supports OPC differentiation, whereas prolonged activation alters lineage outcomes and may impair post-injury repair.

## DISCUSSION

In this study, we investigated the role of Wnt/β-catenin signaling in regulating NG2 glia and their progeny in both healthy and ischemic adult brains. We used transgenic mouse models that enabled lineage tracing of NG2-expressing cells combined with targeted genetic modulation of Wnt pathway activity. Single-cell RNA-seq identified twelve transcriptionally distinct clusters within the OL lineage, with their abundance varying according to ischemic injury and Wnt signaling status. Our data show that canonical Wnt signaling modulates both NG2 glial proliferation and cell fate specification.

We confirmed that FCI activates canonical Wnt signaling in NG2-expressing cells. Although Wnt pathway dysregulation has previously been implicated in ischemia and other neuropathological contexts [25, 67], its cell type-specific roles have not been sufficiently characterized. Using mouse models with either constitutive activation or inhibition of Wnt/β-catenin signaling in NG2 glia, we show that these opposing manipulations do not necessarily produce opposite phenotypic outcomes. This finding suggests that the temporal dynamics and context of Wnt signaling, rather than its simple activation state, are critical determinants of the cellular response. Furthermore, ischemia had a profound impact on gene expression across utilized mouse strains, indicating that the stroke-induced signaling environment exerts dominant effects that may not be fully overridden by genetic modulation of Wnt activity.

Consistent with previous reports, we observed an increased number of NG2-tdTom cells in the lesioned brain region following ischemic injury. Cell proliferation peaked at day 3 post-FCI and remained elevated at day 7, likely reflecting both ongoing proliferation and directed migration toward the lesion site [13]. Although canonical Wnt signaling is generally considered mitogenic, our data suggest this paradigm does not apply to NG2 glia *in vivo*. As shown in Figures 1C and 3B, Wnt hyperactivation resulted in a higher overall proportion of NG2-tdTom cells; however, the fraction of actively proliferating NG2-tdTom cells was reduced in both Wnt-activating and Wnt-inhibiting mouse strains. This challenges the classical view of Wnt as a pro-proliferative signal. One possible explanation is that Wnt signaling promotes a transition from proliferation to differentiation in specific cellular contexts [68]. While Wnt3a has been shown to stimulate NG2 glial proliferation *in vitro* [69], several *in vivo* studies, including a comprehensive analysis by Guo and colleagus [70], report no such effect. Consistent with this, Dkk1-expressing mice showed a substantial reduction in both total NG2 glia and their proliferating subset (Figure 1B,C), likely reflecting enhanced cell cycle arrest (Figure 3C).

To investigate how Wnt signaling influences the various stages of OL maturation, we performed scRNA-seq and analyzed the expression of stage-specific marker genes across all identified clusters. Most cluster-enriched genes corresponded to those previously associated with defined stages of oligodendrogenesis, as summarized in Valihrach and colleagues [40]. However, our clustering was based on a combined dataset of healthy and ischemic cortical tissue. Ischemic samples contained a higher proportion of cells expressing genes linked to OL-associated pathologies, particularly those associated with the MOL-DA (disease-associated mature OL) state (Figure 2D, Supplementary Table S1), likely reflecting injury-induced OL dysfunction or degeneration. Notably, a subset of these cells co-expressed genes characteristic of both OPCs and mature OLs. This hybrid transcriptional profile may indicate lineage plasticity in response to ischemic stress, potentially involving dedifferentiation or state reversion within the OL lineage, as recently described by Bai et al. [71].

Our data show that elevated canonical Wnt signaling impedes OL maturation and myelination. Although β-catenin activity is normally present during OL development, its artificial hyperactivation inhibits (re)myelination [72]. Similarly, mice with constitutive activation of the canonical Wnt pathway display delayed emergence of myelinating OLs [73]. Conversely, inhibition of Wnt signaling via Axin2 stabilization accelerates OL maturation, and the role of Wnt signaling in human oligodendrogenesis has also been documented [74].

The influence of canonical Wnt signaling on OL lineage cells appears highly context- and stage-dependent, as shown during OL development [75–77]. While Wnt activity can inhibit OPC differentiation into OLs, it may enhance myelination at later stages [78]. Thus, precise temporal and spatial regulation of Wnt signaling is critical for proper progression from OPCs to mature, myelinating OLs. Notably, Tcf4, the transcriptional effector of β-catenin, can regulate OL maturation independently of canonical Wnt signaling [79, 80]. This may help explain why Dkk1-mediated Wnt inhibition in NG2 glia did not increase OL numbers, as membrane-level inhibition of Wnt signaling would not affect β-catenin-independent pathways through which Tcf4 might act.

In a recent study, Zhang and colleagues showed that Tcf4 suppresses the expression of bone morphogenetic protein 4 (Bmp4). Overproduction of Tcf4 inhibits autocrine Bmp4 signaling, which impairs the maturation of OPCs [81]. Therefore, it can be speculated that hyperactivation of the Wnt pathway has the opposite effect, resulting in a block of OPC differentiation. Consistent with this, we observed increased *Bmp4* expression in several cell populations isolated from the brains of Ctnnb1(ΔEx3) mice (Supplementary Table S2).

Additionally, re-clustering showed that cluster 2 partially overlapped with the MOL-DA population but also included immature OLs. Along with COP/NFOL cells, this cluster was proportionally most enriched in Ctnnb1(ΔEx3) samples, consistent with a block in OPC differentiation. Notably, a small subset of these “blocked” cells was also found in the SHAM-operated controls, possibly reflecting the elimination of immature OLs during an inefficient or aborted maturation process, as discussed in Hughes and Stockton [82].

Studies investigating NG2 glia subpopulations and their regenerative potential after stroke have consistently highlighted the heterogeneity and plasticity of NG2 glia in the adult brain [11, 15, 83]. We identified transcriptionally distinct NG2 glia subpopulations expressing gene signatures typically associated with astrocytes (“NG2-astro”) and pericytes (“NG2-peri”) in both healthy and ischemic cortex of control mice (Figure 6B and Supplementary Table S4). The ability of NG2 glia to differentiate into astrocytes – demonstrated in both *in vitro* and *in vivo* contexts [84] – appears enhanced under pathological conditions in the CNS [10, 19].

Recently, we described an astrocyte-like NG2 glia subpopulation with features of NPCs that expanded following ischemic stroke [9]. Interestingly, the “NG2-astro” subpopulation identified in the present study expressed astrocytic markers but did not correspond to the previously characterized “astro/NPC-like” subset with progenitor potential. Both populations expressed markers such as *Cldn10* and, post-stroke, *Gfap*. However, the previously described astro/NPC-like subset also expressed NPC-and proliferation-associated genes, including *Ascl1*, *Mcm2*, *Sox11*, and *Pcp4* - genes that, in the current dataset, were restricted to mitotic and basal OPC clusters. This discrepancy may result from differences in dataset structure: the present study combines samples from three transgenic mouse strains, enabling finer clustering resolution, while our previous study included only control strain, comprising a smaller total number of cells.

While the neurogenic potential of NG2 glia remains debated, these cells have several distinctive features that align them with neurons. Developmentally, NG2 glia originating from ventral progenitor domains share a lineage with interneurons [85]. Compared to other glial cell types, such as astrocytes, NG2 glia show reduced levels of repressive histone modifications at key interneuron-associated genes [86]. Collectively, these features position NG2 glia as promising candidates for investigating induced neurogenesis through targeted genetic manipulation.

Our earlier study demonstrated the appearance of neuron-like cells derived from NG2-tdTom cells 28 days after ischemia, which we hypothesized originated from the astro/NPC-like subpopulation, although we lacked specific markers to prospectively identify or lineage-trace these distinct NG2 glial subsets [9]. In the present study, Wnt pathway hyperactivation led to marked expansion of the NG2-astro population and emergence of NG2-derived cells with neuronal morphology in the cortex, even of non-ischemic animals. This observation suggests that NG2-astro cells might serve as a source of these neuron-like cells. However, transcriptomic profiling of NG2-astro cells in Ctnnb1(ΔEx3) mice, when compared to controls, showed enriched embryonic astrocyte and OPC markers. This indicates that sustained Wnt/β-catenin activation reinforces a progenitor-like identity in NG2 glia rather than promoting their differentiation into astrocytes. This effect might be mediated by activation of the Notch signaling pathway (Figure 6E), although, that would somewhat contradicts a recent study by Guo and colleagues showing that Notch signaling promotes astrocyte generation. However, it is important to note that their conclusion was drawn in a different cellular context, specifically during neural stem cell differentiation [87].

Electrophysiological recordings revealed that NG2-derived cells with neuronal morphology exhibited AP firing (Figure5). While generation of fast, voltage-dependent Na^+^ currents in NG2 glia has been reported, it typically does not meet the criteria of neuronal APs [88–90]. In contrast, cells from the somatosensory cortex of our Ctnnb1(ΔEx3) mice exhibited action potentials resembling those of neurons, indicating changes in membrane properties, including a higher density of voltage-gated Na^+^ channels (Supplementary Table S3). Interestingly, cortical NG2 glia from SHAM animals of all three mouse strains (Ctnnb1(ΔEx3), Dkk1, control strain) did not show any Na^+^ currents, in contrast to earlier studies conducted in different brain regions (e.g., hippocampus [89]) or at different ontogenetic stages of animals used (e.g., postnatal day 33 [88]).

Collectively, our findings suggest that overactivation of Wnt signaling may induce neuronal differentiation from NG2 glia in a region-specific manner. As described in the Results section, hyperactivation of the Wnt/β-catenin pathway maintains COP/NFOL cells in a progenitor-like state, suggesting that some of these cells could potentially change their identity. However, the genetic modification in Ctnnb1(ΔEx3) mice results in constitutive Wnt/β-catenin pathway activation, which does not fully replicate the physiological conditions in wild-type mice after ischemia. For future translational considerations, it will be important to further examine these findings in additional experimental settings. This may include analyses in both sexes, the use of complementary ischemia models, and evaluation of clinically relevant Wnt modulators such as LiCl. Extending these observations to human-derived cells or tissues would also help to assess the relevance of Wnt-dependent regulation of NG2 glia plasticity for post-stroke repair.

## MATERIAL AND METHODS

### Transgenic animals

All procedures involving animals used for scientific purposes were performed in accordance with the Directive of the European Parliament and the Council (2010/63/EU) and the Animal Welfare Guidelines approved by the animal-welfare body of the Institute of Experimental Medicine of the Czech Academy of Sciences. The following mouse strains were used in this study: Rosa26-tdTomato [B6;129S6-Gt(ROSA)26Sor^tm14(CAG-tdTomato)Hze/J^], NG2CreER^TM^BAC [B6.Cg-Tg(Cspg4-cre/Esr1*)BAkik/J], and Pdgfrα^EGFP^ [B6.129S4-Pdgfra^tm11(EGFP)Sor/J^], all obtained from The Jackson Laboratory (JAX stock #007908, #008538 and #007669, respectively). Rosa26-Dkk1 mice expressing the extracellular Wnt pathway inhibitor Dkk1 following Cre-mediated excision of a transcriptional blocker [31] were kindly provided by Dr. Long (Children’s Hospital of Philadelphia, USA). Ctnnb1^lox(ex3)^ mice carrying a conditional (“floxed”) allele of the *Ctnnb1* gene encoding β-catenin [32] were provided by Dr. Taketo (Kyoto University, Japan). In this model, Cre-mediated recombination leads to stabilization of the β-catenin protein. Young adult male mice (P59–P69 at the beginning of the experiment) were used to avoid the potential neuroprotective effects of estrogen in female animals [91]. DNA recombination was induced by two consecutive intraperitoneal injections of tamoxifen (200 mg/kg body weight; # T006000, Toronto Research Chemicals) dissolved in corn oil (#C8267, Sigma-Aldrich). Experimental treatments (see below) started 15 days after the second tamoxifen injection.

### Induction of FCI

Focal cerebral ischemia was induced using a well-established permanent distal MCAO model [92]. The entire surgical procedure has been described in detail previously [25]. In sham-operated control animals, the same procedure, including craniotomy, was performed, but the MCA was not occluded. This MCAO model is characterized by a high survival rate (>95%) and good reproducibility and usually results in a small, well-demarcated infarct localized mainly in the cortical area [93]. The animals were examined 1, 3, 7, 12, and 28 days post-surgery. At least two animals per group (animals after MCAO or sham-operated) were examined for each time point.

### Tissue isolation and immunohistochemistry

All animals were deeply anesthetized with pentobarbital (#Y0002194, 100 mg/kg, Sigma-Aldrich) prior to tissue collection. Transcardial perfusion with saline containing heparin and consequent fixation with 1% paraformaldehyde was performed; detailed procedures for preparing tissue sections for histology and immunofluorescence staining have been described previously [25]. The following primary antibodies were used: goat polyclonal anti-ChAT (1:100, #AB144P, Millipore), mouse monoclonal anti-GABA (1:300, #A2052, Sigma-Aldrich), mouse monoclonal anti-GFAP conjugated to Alexa Fluor 488 (1:300; clone GA5; #53-9892, eBioscience), mouse monoclonal anti-Mag (1:800; clone 513; #MAB1567, Millipore), mouse monoclonal anti-Map2 (1:800, #MAB3418, Chemicon), mouse monoclonal anti-NeuN (1:100; clone A60; #MAB377, Chemicon), goat polyclonal anti-Pdgfrα (1:500; #AF1062, R&D Systems), mouse monoclonal anti-Pvalb (1:500, #P3088, Sigma-Aldrich). Secondary antibodies included: donkey polyclonal anti-mouse IgG conjugated to Alexa Fluor 488 (1:200; #A32766, Thermo Fisher Scientific), donkey polyclonal anti-goat IgG conjugated to Alexa Fluor 647 (1:200; #A32849, Thermo Fisher Scientific). Cell nuclei were counterstained with 300 nM 4′,6-diamidino-2-phenylindole (DAPI; #202710100, Thermo Fisher Scientific).

### Preparation of single-cell suspension and FACS

Single-cell suspensions were prepared according to a previously described protocol [9]. The dorsal and dorsolateral cortex of the ischemia affected hemisphere (and the corresponding region in sham-operated animals) was excised from approximately 600 µm thick coronal brain slices. The cortical tissue was cut into small pieces and incubated for 40 minutes at 37 °C with constant shaking in 1 ml papain solution (20 U/ml; #LK003176, Worthington Biochemical Corporation) prepared in an isolation buffer supplemented with DNase I (120 U/ml; #LK003170, Worthington). During enzymatic digestion, the tissue was gently triturated every 15 minutes with a wide bore 1-ml pipette tip to facilitate mechanical dissociation. The enzymatic reaction was terminated by adding an ovomucoid protease inhibitor solution (0.95 mg/ml; #LK003182, Worthington) in Earle’s balanced salt solution (EBSS; #LK003188, Worthington) supplemented with DNase I (95.2 U/ml). The resulting cell suspension was further purified with Debris Removal Solution (#130-109-398, Miltenyi Biotec) according to the manufacturer’s instructions. The cells were then incubated at 4 °C for 15 minutes with antibodies against CD45 (#102422, BioLegend) and CD31 (#103126, BioLegend) to label lymphocytes and endothelial cells, respectively. The antibodies were diluted in Neurobasal-A medium (#10888-022, Life Technologies) supplemented with 2% B27 (#17504-044, Life Technologies). After staining, the cells were washed and resuspended in Neurobasal-A medium at 4 °C.

Viable cells identified as Hoechst 33258-negative (#H1398, Life Technologies) and CD45/CD31-negative were isolated by FACS (BD Influx). Sorted cells were collected in 1.5 ml Eppendorf tubes containing 200 µl Neurobasal-A medium with 2% B27. These cells were processed either for scRNA-seq sequencing using the 10x Genomics platform or isolated using the RNeasy Micro Kit (#74004, Qiagen).

### Reverse-transcription quantitative polymerase chain reaction (RT-qPCR)

Complementary DNA was synthesized using random hexamer primers and Maxima Reverse Transcriptase (200 U/µL; #EP0741, Thermo Fisher Scientific) according to the manufacturer’s protocol. PCR reactions were performed in triplicate using the LightCycler® 480 SYBR Green I Master Mix (#04887352001, Roche Diagnostics) in a LightCycler^®^ 480 instrument (Roche Diagnostics). The primer sequences used for RT-qPCR are listed in Supplementary Table S6.

### Single-cell RNA-seq

Barcoded single-cell cDNA libraries were prepared using the Chromium Controller (10x Genomics) and the Chromium Next GEM Single Cell 3′ Reagent Kit v3.1 according to the manufacturer’s protocol. The resulting libraries were pooled and sequenced on an Illumina NextSeq 500 platform, achieving a sequencing depth of over 100,000 reads per cell. The generated datasets are available in the ArrayExpress repository in the EMBL-EBI BioStudies database under the following accession numbers: E-MTAB-14958: Constitutive activation of Wnt/β-catenin signaling; E-MTAB-14905: Inhibition of Wnt signaling; E-MTAB-11967: Control strain (previously published in [9]. Data analysis was performed using RStudio (RStudio, PBC) and Seurat package version 4 [94]. All datasets were merged into a combined dataset of 18,951 cells. Cell clustering was performed using the Louvain algorithm, which is based on principal component analysis (PCA). Uniform Manifold Approximation and Projection (UMAP) was used for nonlinear dimensionality reduction to visualize the data and confirm the cluster assignment [95]. The identified cell clusters were further analyzed based on the expression of specific marker genes using a molecular atlas of adult brain cells [96]. Cells expressing markers indicative of perivascular cells and microglia were excluded from further analyses. The final dataset comprised 4,781 cells of the OL lineage. Pseudotime trajectory analysis was done using Monocle3 [62]. GSEA was performed using clusterProfiler version 4.12.6 (https://code.bioconductor.org/browse/clusterProfiler/) and org.Mm.eg.db 3.19.1 (https://bioconductor.org/packages/org.Hs.eg.db/). ORA was performed using DEGs (with adjusted p-value < 0.05 and |log2 FC | ≥ 1 as significance criterions) by the Enrichr platform [36, 37].

### Patch-clamp recording

Preparation of coronal brain slices and electrophysiological measurements were performed as previously described [9]. In brief, anesthetized animals were perfused transcardially with ice-cold N-methyl-D-glucamine (NMDG)-containing isolation solution. Coronal sections (200 µm thick) were incubated for 30 minutes at 34 °C in NMDG isolation solution and for another 30 minutes in artificial cerebrospinal fluid (aCSF). To investigate the membrane properties of NG2 glia and their progeny, the patch-clamp technique was used in the whole-cell configuration. Pre-incubated tissue sections were kept in the recording chamber at room temperature, perfused with aCSF and gassed with 95% O_2_ and 5% CO_2_. Recording pipettes with a tip resistance of 8-12 MΩ were made of borosilicate capillaries (#BF150-86-10, Sutter Instruments) and filled with intracellular solution. A more detailed description of the patch-clamp technique, including the composition of the solution, can be found in our previous publications [9, 25]. Membrane properties of NG2 glia were recorded 3 days after surgery in both SHAM and MCAO animals; tdTomato-positive OLs and perivascular cells were excluded from the analysis based on the following parameters. Pericytes were identified as elongated tdTomato-positive cells near blood vessels showing negligible inward and outward K^+^ currents; OLs were identified by their typical current pattern, i.e., decaying time- and voltage-independent K^+^ currents with distinct tail currents [97, 98].

The electrophysiological data were measured at a sampling frequency of 10 kHz with an EPC9 or an EPC10 amplifier, controlled by the PatchMaster software (HEKA Elektronik) and filtered with a Bessel filter. V_M_ was measured by switching the amplifier to current clamp mode. Using FitMaster software (HEKA Elektronik), IR was calculated from the current value 40 ms after the start of the depolarizing 10 mV pulse from the holding potential of -70 mV. C_M_ was calculated from the area-under-curve of a first current transient induced by a 10 pA hyperpolarizing pulse, as described in a publication by Golowasch and colleagues [99]. This approach replaced assessing the C_M_ value from the lock-in protocol so that the values are comparable to recordings obtained from the MultiClamp 700 B amplifier.

The recording protocols as well as the steps to isolate specific K^+^ currents were the same as in our previous publications [9, 25]. The current densities were calculated by dividing the maximum current amplitudes by the corresponding C_M_ values for each cell.

To capture the electrophysiological properties of cells with neuron-like morphology, the tissue isolation protocol and aCSF solutions were adopted from [100]. Transcardial perfusion was performed with a 4 °C cold neuroprotective isolation solution containing (in mM): 92 (NMDG)-Cl, 2.5 KCl, 1.25 NaH_2_PO_4_, 30 NaHCO_3_, 20 HEPES, 25 glucose, 2 thiourea, 5 L-ascorbic acid sodium salt, 3 pyruvic acid sodium salt, 0.5 CaCl_2_, and 10 MgCl_2_ (pH 7.4); osmolality 305 ± 5 mOsmol/kg. Coronal slices (300 µm thick) were incubated for 12 minutes in a 30 °C neuroprotective isolation solution and then transferred to aCSF solution at room temperature containing (in mM): 92 NaCl, 2.5 KCl, 1.25 NaH_2_PO_4_, 30 NaHCO_3_, 20 HEPES, 25 glucose, 2 thiourea, 5 L-ascorbic acid sodium salt, 3 pyruvic acid sodium salt, 2 CaCl_2_, and 7 MgCl_2_; osmolality 305 ± 5 mOsmol/kg. Tissue sections were placed in the recording chamber at room temperature and perfused with aCSF containing (in mM) 119 NaCl, 2.5 KCl, 1.25 NaH_2_PO_4_, 24 NaHCO_3_, 12.5 glucose, 2 CaCl_2_, and 2 MgCl_2_; osmolality 305 ± 5 mOsmol/kg. All solutions were gassed with 95% O_2_ and 5% CO_2_. Recording pipettes with a tip resistance of 5 ± 2 MΩ were prepared from borosilicate capillaries and filled with intracellular solution containing (in mM): 107.5 K-gluconate, 32.5 KCl, 10 HEPES, 5 EGTA, 4 ATP magnesium salt, 0.6 GTP sodium salt, and 10 phosphocreatine di(tris) salt (pH 7.2); osmolality 310 ± 5. To measure the spike activity of neuron-like cells, depolarization pulses from 0 pA to 1000 pA were applied in 100 pA steps in current-clamp mode. For this experiment, the EPC10 amplifier controlled by the PatchMaster software or a MultiClamp 700 B amplifier in combination with the AxonTM DigiData 1550 B digitizer controlled by the Clampex software (Axon Instruments) were used.

### Confocal microscopy and image analysis

Localization of cell types observed throughout the cortical area surrounding the ischemic injury was assessed using a Dragonfly confocal fluorescence microscope with a 530 Andor rotating disk (Oxford Instruments) equipped with a Zyla 4.2 PLUS sCMOS camera and Fusion acquisition system. Tiled images were obtained by superimposing 50-70 individual confocal images acquired with a 20× objective. The image series were then digitally fused using the Fusion Stitching Tool. The acquired images were processed using Imaris visualization software (Oxford Instruments). Co-localization of the antibody immunofluorescence signal and endogenous fluorescence of NG2 glia was examined using the Leica Stellaris confocal platform (Leica Microsystems). The immunopositive areas of the cortex surrounding the ischemic lesion were quantified using Fiji ImageJ software (National Institutes of Health) [101]. For each quantification, a total of six cortical sections from at least two biological replicates were analyzed.

### Data analysis and statistics

Data are presented as mean ± standard error of the mean (SEM) or mean ± standard deviation (SD) for a given number (n) of samples/cells, as indicated in each figure. Repeated-measures ANOVA was performed to detect significant differences between experimental groups in immunofluorescence quantification; one-way ANOVA was used to detect significance in RT-qPCR. The electrophysiological properties were then compared with the Kruskal-Wallis post-test using the one-way ANOVA. Values of p < 0.05 were considered significant. The normality distribution of the data was checked for each set of measurements.

## DATA AVAILIBILITY STATEMENT

The raw data that support the findings of this study are openly available in the ArrayExpress database (https://www.ebi.ac.uk/biostudies/arrayexpress) under the following accession numbers: E-MTAB-11967, E-MTAB-14905, and E-MTAB-14958.

## ACKNOWLEDGEMENTS

We thank Sarka Takacova for critical reading of the manuscript and Helena Pavlikova, Sarka Kocourkova, and Zdenek Cimburek for their technical assistance. We are grateful to Dr. Kozmik for providing the Ctnnb1^lox(ex3)^ mice. This research was funded by the Czech Science Foundation, grant no. 24-10912S, and by the Czech Academy of Sciences (Strategy AV21, grant no. VP29). TK was in part supported by the Grant Agency of Charles University (grant no. 244223). MK, JKu and KV were in part supported by the Ministry of Education, Youth and Sports (ELIXIR CZ - LM2023055). The Light Microscopy Core Facility, IMG CAS, Prague, Czech Republic, was supported by the Ministry of Education, Youth and Sports (LM2023050, CZ.02.1.01/0.0/0.0/18_046/0016045 and CZ.02.01.01/00/23_015/0008205).

## AUTHOR CONTRIBUTIONS

MA, VK, and LJ designed the study concept and supervised the project. MA, VK, TK, and LJ coordinated the experiments. TK, JKr, and AF were responsible for the tissue isolation and processing; TK and DK performed MCAO surgeries. JKu, KV, and MK carried out bioinformatics processing; LJ facilitated analysis of the gene expression data from scRNA-seq and RT-qPCR experiments. TK and JKr performed the electrophysiological study. TK and LJ acquired and analyzed data from the immunochemical experiments. LJ supervised the breeding of the transgenic mouse strains. TK and LJ wrote the manuscript. JKr, VK, and MA revised the manuscript. MMT provided the mouse strain crucial for the manuscript. All authors read and approved the final version of the manuscript.

## ABRREVIATIONS

Abca2: ATP-binding cassette sub-family A member 2
Abcc9: ATP-binding cassette sub-family C member 9
aCSF: Artificial cerebrospinal fluid
Aif1: Allograft inflammatory factor 1
Akt: Protein kinase B
Anxa2: Annexin A2
AP: Action potential
Ascl1: Achaete-scute family bHLH transcription factor 1
BBB: Blood-brain barrier
Bub1: Mitotic checkpoint serine/threonine kinase
Calb2: Calbindin 2
Ccna2: Cyclin A2
Ccnd1: Cyclin D1
Cdkn1a/c: Cyclin-dependent kinase inhibitor 1A/C
Cdkn2a: Cyclin-dependent kinase inhibitor 2A
ChAT: Choline acetyltransferase
Cldn11: Claudin 11
C_M_: Membrane capacitance
Cnp: 2′,3′-cyclic-nucleotide 3’-phosphodiesterase
CNS: Central nervous system
COP: Committed oligodendrocyte precursor
Ctnnb1: Catenin, beta 1
Csfr: Colony-stimulating factor receptor
CTX: Cortex
Dcx: Doublecortin
DEG: Differentially expressed gene
Dkk1: Dickkopf-1
EBSS: Earle’s balanced salt solution
Emp3: Epithelial membrane protein 3
ER: Estrogen receptor
Ex3: Exon 3
FACS: Fluorescence-activated cell sorting
FBS: Fetal bovine serum
FCI: Focal cerebral ischemia
Foxo1: Forkhead box protein O1
GABA: Gamma-aminobutyric acid
GFAP: Glial fibrillary acidic protein
Gjb1: Gap junction protein, beta 1
Gpr17: G-protein coupled receptor 17
Gria2: Glutamate ionotropic receptor AMPA-type subunit 2
GSEA: Gene set enrichment analysis
Hcn2: Hyperpolarization-activated cyclic nucleotide-gated ion channel 2
Hes1: Hairy and enhancer of split 1
Hey2: Hairy/enhancer of split-related with YRPW motif 2
IR: Input resistance
K_A_: A-type potassium current
Kcnj8: Potassium inwardly rectifying channel subfamily J member 8
K_DR_: Delayed rectified potassium current
K_IR_: Inwardly rectified potassium current
Klk6: Kallikrein-related peptidase 6
Mag: Myelin-associated glycoprotein
Mal: Myelin and lymphocyte protein
Map2: Microtubule-associated protein 2
MCAO: Middle cerebral artery occlusion
Mcm2: Minichromosome maintenance complex component 2
MFOL: Myelin-forming oligodendrocyte
Mki67: Marker of proliferation Ki-67
Mobp: Myelin-associated oligodendrocyte basic protein
Mog: Myelin oligodendrocyte glycoprotein
MOL: Mature oligodendrocyte
MOL-DA: Disease-associated mature oligodendrocyte
NCSs: Neural stem cells
Myrf: Myelin-gene regulatory factor
NeuN: Neuronal nuclei (Rbfox3 gene product)
NG2: Neuron-glial antigen 2
Nkd1: Naked cuticle 1
NMDG: N-methyl-D-glucamine
NPC: Neural progenitor cell
NSC: Neural stem cell
OL: Oligodendrocyte
Olig1/2: Oligodendrocyte transcription factors 1 and 2
Opalin: Oligodendrocytic myelin paranodal and inner loop protein
OPC: Oligodendrocyte precursor cell
ORA: Over-representation analysis
PCA: Principal component analysis
PCNA: Proliferating cell nuclear antigen
Pdgfrα: Platelet-derived growth factor receptor alpha
Plk1: Polo-like kinase 1
Plvap: Plasmalemma vesicle-associated protein
Pvalb: Parvalbumin
Rbfox3: RNA-binding fox-1 homolog 2
RT-qPCR: Reverse transcription quantitative polymerase chain reaction
SD: Standard deviation
SEM: Standard error of the mean
scRNA-seq: Single-cell RNA sequencing
Serpina3n: Serine (or cysteine) peptidase inhibitor, clade A, member 3N
Sox2: SRY-box transcription factor 2
Spc24/25: Spindle pole component 24/25
Sp5: Specificity protein 5
ST: Striatum
SVZ: Subventricular zone
Syp: Synaptophysin
Tcf4: T-cell factor 4 (protein)
Tcf7l2: Transcription factor 7-like 2 (gene)
Timp3: Tissue inhibitor of metalloproteinase 3
Tnfrsf19: Tumor necrosis factor receptor superfamily member 19
Tpx2: Microtubule nucleation factor
TTX: Tetrodotoxin
Ube2c: Ubiquitin conjugating enzyme E2 C
UMAP: Uniform manifold approximation and projection
V_M_: Resting membrane potential

