## Supplementary material for "Canonical Wnt signaling controls the fate and plasticity of NG2 glia in the healthy and ischemic adult mouse cortex": Supplementary Material.docx

**Supplementary Figure and Table Legends**

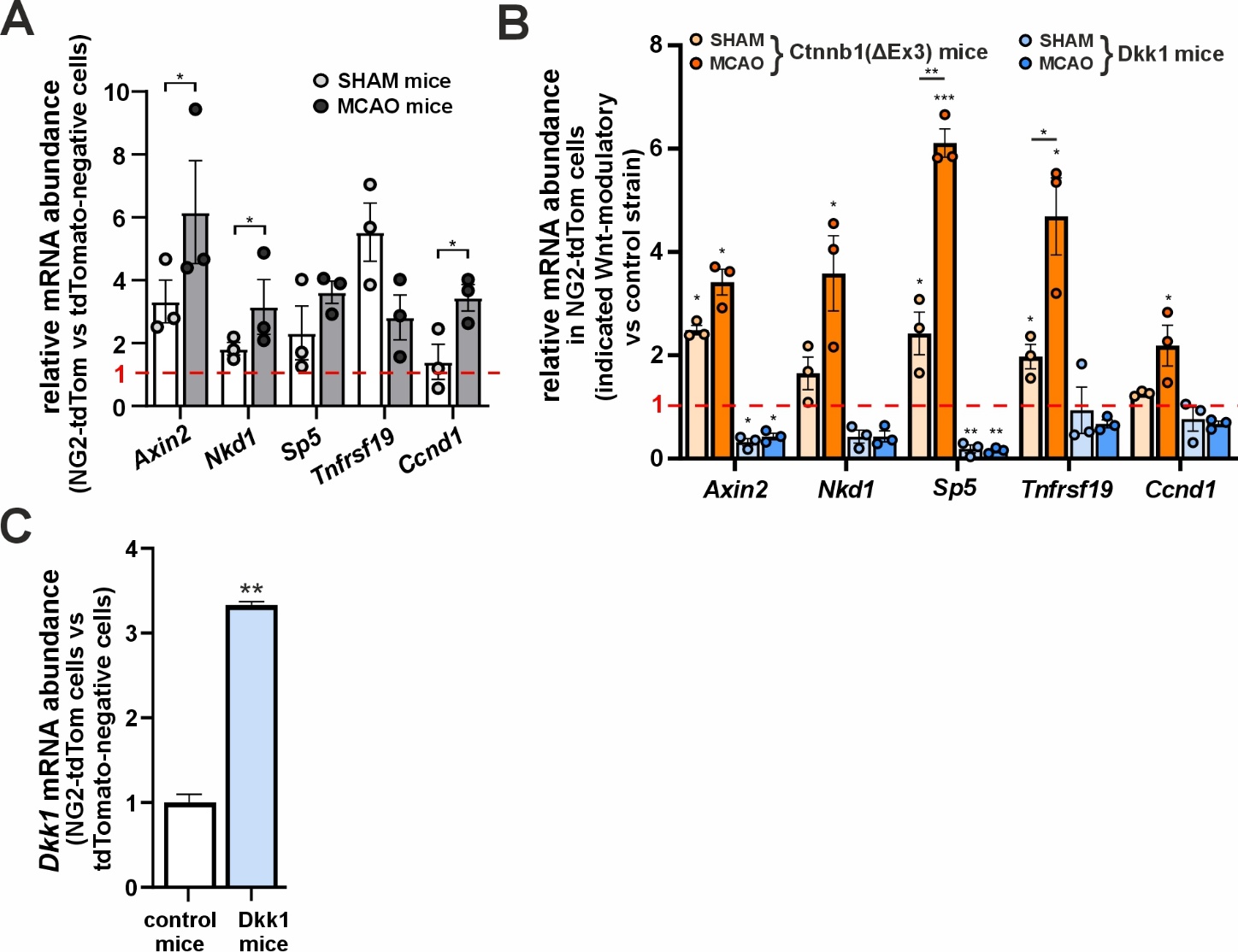

**Supplementary Figure S1.** **Validation of the Wnt signaling level in NG2-expressing cells.** NG2^+^ cells with tamoxifen-induced production of tdTomato red fluorescent protein (NG2-tdTom) were isolated using fluorescent-activated cell sorting from the mouse cortex. Gene expression was analyzed using reverse transcription followed by quantitative polymerase chain reaction (RT-qPCR).

(A) Analysis of Wnt/β-catenin target gene relative expression (*Axin2*, *Nkd1*, *Sp5*, *Tnfrsf19*, and *Ccnd1*) in NG2-tdTom cells sorted from the cortex of control mice. Middle cerebral artery occlusion (MCAO) or sham operation (SHAM) were performed 3 days before the NG2-tdTom cell isolation. Expression was normalized to tdTomato-negative cells from the same animals (set to 1) and to the reference gene *Ubb*. Data are means ± SD of technical triplicates of 5,000 cells per group.

(B) Relative expression of the Wnt responsive genes in Wnt modulatory strains was inspected at day 3 post-surgery, i.e., 18 days after the tamoxifen-mediated activation of the Wnt modulatory allele. Gene expression of the corresponding gene in the control strain was arbitrarily set to 1.

(C) Quantification of the Dickkopf 1 (*Dkk1*) Wnt inhibitor expression NG2-tdTom cells in comparison to other cells isolated from the cortex. Statistical significances were determined using the one-way ANOVA test; *, p < 0.05; **, p < 0.01. Ccnd1, cyclin D1; Ctnnb1, catenin beta 1; ΔEx3, deleted exon3; Nkd1, naked cuticle 1; Sp5, transcription factor Sp5; Tnfrsf19, tumor necrosis factor receptor superfamily member 19.

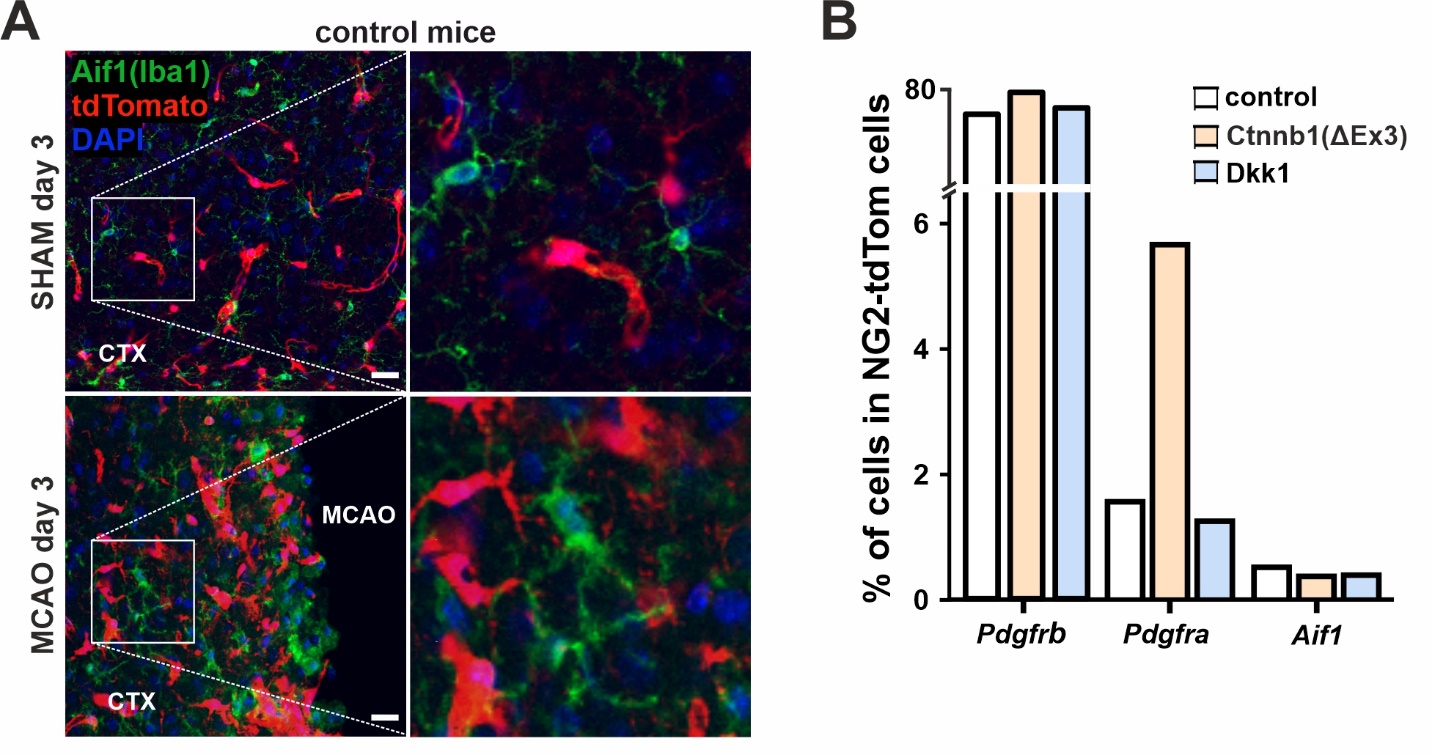

**Supplementary Figure S2. Wnt signaling modulation mostly influences NG2 glia within NG2-tdTom cells in the mouse cortex (CTX).** Single cell RNA-sequencing (scRNA-seq) analysis of NG2-tdTom cells from the ischemic (MCAO) and healthy (SHAM) cortex of the three analyzed mouse strains. A pooled dataset contained 18,951 cells.

(A) Representative slides show staining of NG2-tdTom cells and microglia (labeled with Aif1/Iba1 protein, green) in the healthy and ischemic cortex (CTX) 3 days after the surgery. Insets are shown on the right side. Samples were counterstained with DAPI nuclear stain; scale bars correspond to 25 μm.

(B) Column graph showing the percentage of *Pdgfrb*^+^ (perivascular cells), *Pdgfra*^+^ (NG2 glia), and *Aif1*^+^ (microglia) cells within the SHAM datasets of the respective mouse strains. While the abundance of *Pdgfra*^+^ NG2 glia varies depending on the Wnt pathway modulation, the abundance of *Pdgfrb*^+^ perivascular cells and *Aif1*^+^ microglia remains similar across strains. *Aif1*, allograft inflammatory factor 1; *Pdgfra/b*, platelet-derived growth factor receptor alpha/beta.

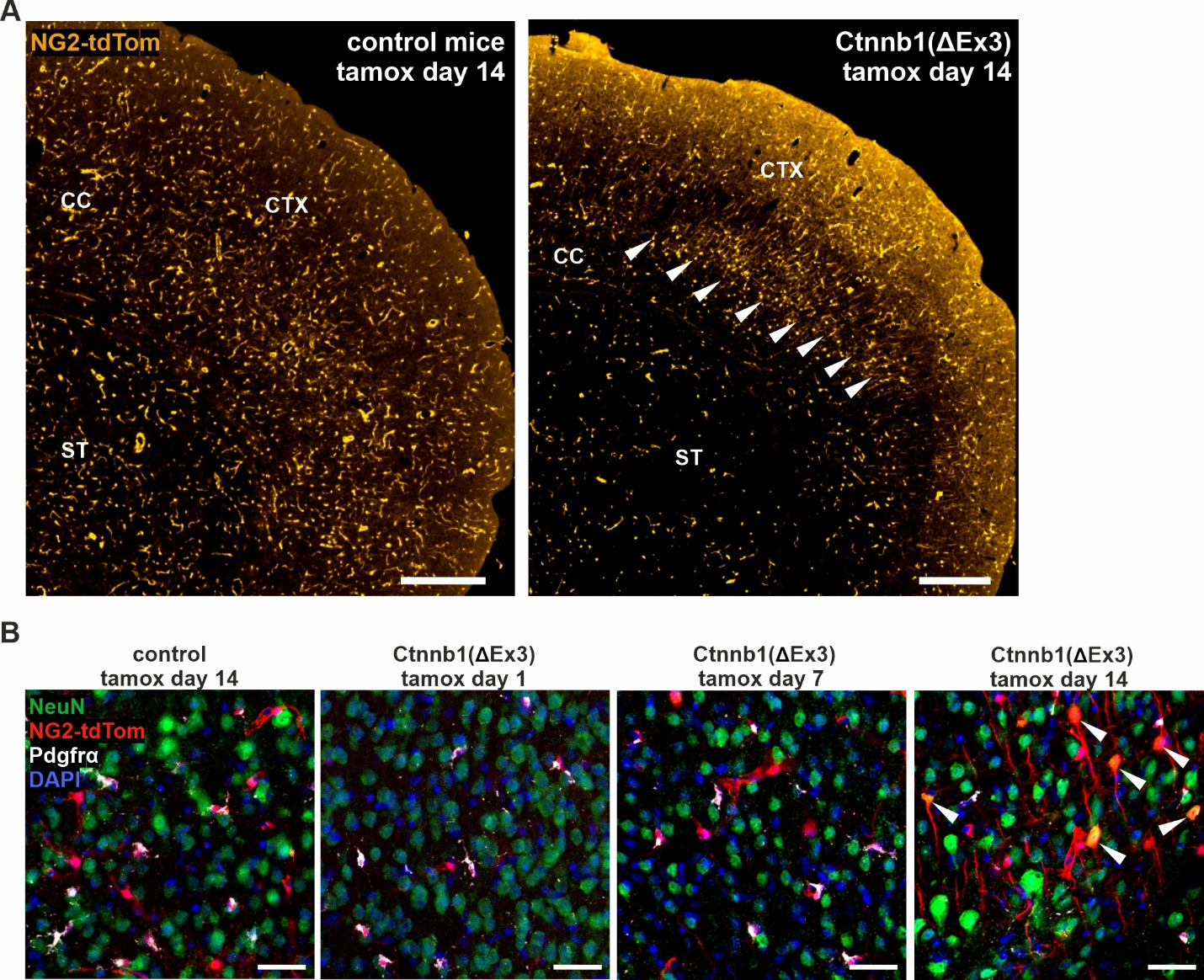

**Supplementary Figure S3. NeuN-positive NG2-tdTom cells are present in the cortex 14 days after the Wnt/β-catenin pathway upregulation.** Mice were treated with tamoxifen for two consecutive days and the brains were examined 1, 7, and 14 days afterwards.

(A) Macrographs showing endogenous fluorescence of NG2-tdTom cells in the striatum (ST), corpus callosum (CC), and cortex (CTX) 14 days after the second tamoxifen injection. Note the increased concentration of NG2-tdTom cells with long straight protrusions cumulated in the part of CTX close to CC by the mice with constitutively active β-catenin (white arrowheads). Scale bars correspond to 300 μm.

(B) Fluorescent images of CTX showing that most NG2-tdTom cells in the control strain and Ctnnb1(ΔEx3) strain one and seven days after tamoxifen treatment were positive for NG2 glia marker Pdgfrα (white signal). In contrast, we noted NeuN-positive Ex3/NG2-tdTom cells 14 days after tamoxifen treatment (white arrowheads). Scale bar: 50 μm.

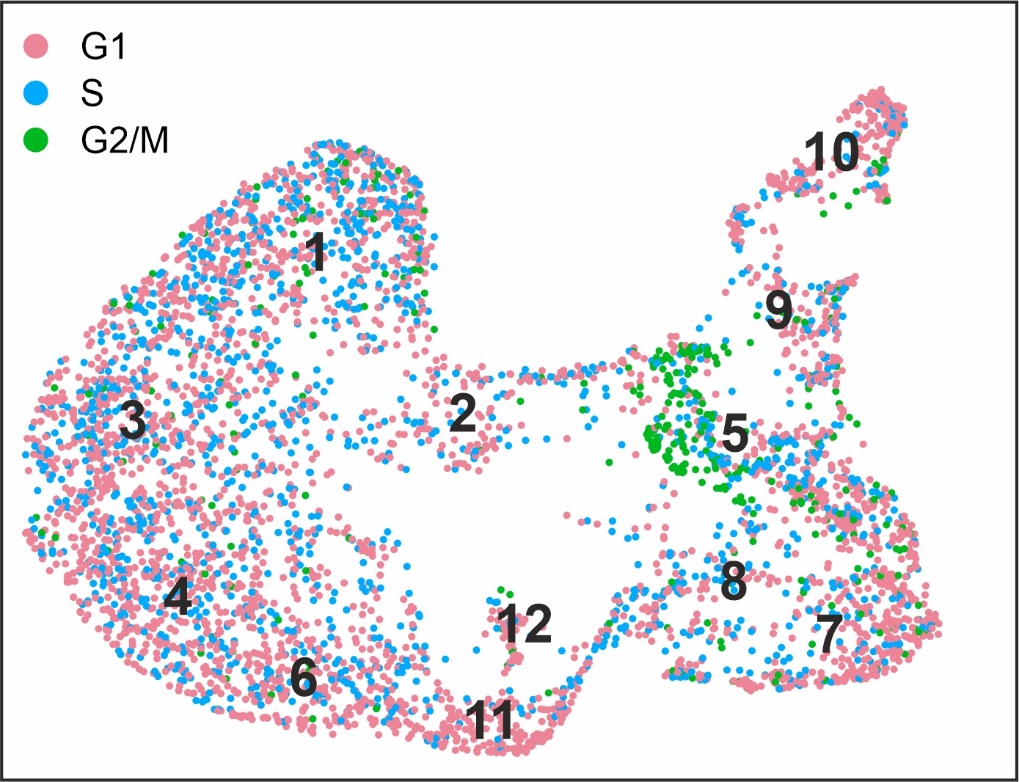

**Supplementary Figure S4. Cell cycle analysis in clusters identified by scRNA-seq within the oligodendroglial (OL) lineage of NG2-tdTom cells.** The diagram shows a UMAP (Uniform Manifold Approximation and Projection) plot colored according to the cell cycle phase. Positions of the 12 identified cell clusters are indicated by their numbers.

**Supplementary Table S1**

Marker genes specifically expressed in each cluster identified within NG2-tdTom cells from the merged dataset of SHAM and MCAO samples (control strain; UMAP shown in Figure 2A) are listed in the corresponding sheet. Genes were considered significantly enriched if they showed average expression |log_2_ FC| > 0.25 and p-value < 0.05 (adjusted using false discovery rate (FDR) correction based on the total gene count in the dataset). For each gene, the percentage of expressing cells in the first and second most enriched clusters is provided. COP, committed oligodendrocyte progenitor cells; NFOL, newly formed oligodendrocytes; MFOL, myelin forming oligodendrocytes; MOL, mature oligodendrocytes; MOL-DA, mature oligodendrocytes disease-associated.

**Supplementary Table S2**

Differentially expressed genes (DEGs) in the OL lineage clusters (Figure 2) isolated from MCAO samples of mice with Wnt signaling modulation compared to the corresponding sample from the control strain (CTRL). The percentage of cells with the gene expression in the first and second analyzed clusters is indicated. Genes with an adjusted p-value < 0.05, UMIs > 50, and pct of expressing cells in the cluster of both samples > 0 are shown. False discovery rate (FDR) correction based on the total number of genes in the dataset was used to adjust the p-value.

**Supplementary Table S3**

Electrophysiological properties of cortical NG2 glia were compared across three mouse strains (control, Dkk1, Ctnnb1(ΔEx3)) and two treatments (SHAM vs. MCAO D3). Additionally, the table includes electrophysiological properties of cells from the somatosensory cortex exhibiting axon-like processes, compared with cortical NG2 glia from sham-operated control mice and the Ctnnb1(ΔEx3) mice (“WNT ON”). Data are presented as mean ± S.E.M. Non-parametric Kruskal-Wallis ANOVA was used and correction for multiple comparison was performed in GraphPad Prism software (v. 8.4.3).

**Supplementary Table S4**

Genes specifically expressed in each identified subpopulation of the OL lineage from both SHAM and MCAO samples isolated from the control strain (UMAP shown in Figure 5A) are listed in the corresponding sheet. Genes with the average expression |log_2_ FC| > 0.25 and p-value < 0.05 were considered as significant. FDR correction based on the total number of genes in the dataset was used to adjust the p-value. For each gene, the table includes log₂ fold change, p-value, FDR-adjusted p-value, and the percentage of cells expressing the gene in the first and second most significantly enriched clusters.

**Supplementary Table S5**

DEGs in the OL lineage subpopulations isolated from MCAO samples of mice with Wnt signaling modulation compared to the corresponding sample from the control strain (CTRL). The percentage of cells with the gene expression in the first and second analyzed clusters is indicated. Genes with an adjusted p-value < 0.05, UMIs > 50, and pct of expressing cells in the cluster of both samples > 0 are shown. FDR correction based on the total number of genes in the dataset was used to adjust the p-value.

**Supplementary Table S6**

Sequences of primers used in the RT-qPCR experiments.
